# Neuronal growth regulator 1 (NEGR1) promotes synaptic targeting of glutamic acid decarboxylase 65 (GAD65)

**DOI:** 10.1101/2022.02.08.479601

**Authors:** Rui Ping Amanda Tan, Irina Kozlova, Feifei Su, Saroj Sah, Ryan Keable, D. Walker Hagan, Sonia Bustamante, Ximing Du, Brenna Osborne, Nigel Turner, Kelly J. Clemens, Denovan Begg, Edward A. Phelps, Hongyuan Yang, Iryna Leshchyns’ka, Vladimir Sytnyk

## Abstract

Neuronal growth regulator 1 (NEGR1) is a glycosylphosphatidylinositol-anchored cell adhesion molecule encoded by an obesity susceptibility gene. We demonstrate that NEGR1 accumulates in GABAergic inhibitory synapses in hypothalamic neurons, a GABA-synthesizing enzyme GAD65 attaches to the plasma membrane, and NEGR1 promotes clustering of GAD65 at the synaptic plasma membrane. GAD65 is removed from the plasma membrane with newly formed vesicles. The association of GAD65 with vesicles results in increased GABA synthesis. In NEGR1 deficient mice, the synaptic targeting of GAD65 is decreased, the GABAergic synapse densities are reduced, and the reinforcing effects of food rewards are blunted. In mice fed a high fat diet, levels of NEGR1 are increased and GAD65 abnormally accumulates at the synaptic plasma membrane. Our results indicate that NEGR1 regulates a previously unknown step required for synaptic targeting and functioning of GAD65, which can be affected by bidirectional changes in NEGR1 levels causing disruptions in the GABAergic signaling controlling feeding behavior.

## Introduction

Neuronal growth regulator 1 (NEGR1) is a cell adhesion molecule of the IgLON family of the immunoglobulin (Ig) superfamily. Members of the IgLON family, which also includes the opioid-binding cell adhesion molecule, neurotrimin, limbic system-associated membrane protein and IgLON family member 5, comprise three extracellular Ig-like domains attached to the plasma membrane (PM) by a glycosylphosphatidylinositol (GPI) anchor (Funatsu et al., 1999; Miyata et al., 2000; Tan et al., 2017). NEGR1 is broadly expressed in the brain (Lee et al., 2012) and accumulates in synapses (Hashimoto et al., 2008; Miyata et al., 2003). NEGR1 molecules bind to each other thereby forming homophilic adhesive bonds connecting synaptic membranes (Miyata et al., 2003; Ranaivoson et al., 2019; Venkannagari et al., 2020). NEGR1 heterophilically interacts with other IgLONs (Miyata et al., 2003; Ranaivoson et al., 2019) and the fibroblast growth factor receptor 2 (FGFR2) (Szczurkowska et al., 2018).

The *NEGR1* gene is associated with obesity in humans (Locke et al., 2015; Speliotes et al., 2010; Thorleifsson et al., 2009). Deletions of the *NEGR1* gene are found in patients with developmental co-ordination disorder, attention deficit / hyperactivity disorder, learning disability, delayed speech and language development, and dyslexia (Genovese et al., 2015; Tassano et al., 2015; Veerappa et al., 2013). Loss of NEGR1 in the arcuate nucleus (ARC) and ventromedial hypothalamus (VMH) of rats causes an increase in food intake (Boender et al., 2014). NEGR1 knock-out (NEGR1-/-) mice show reduced food consumption and body weight (Lee et al., 2012), increased anxiety and depression-like behaviors (Noh et al., 2019), and impaired social behavior (Singh et al., 2018). Increased susceptibility to pentylenetetrazol-induced seizures of these animals (Singh et al., 2018) suggests that NEGR1 is involved in control of neuronal activity via currently unknown mechanisms.

Inhibitory γ-amino butyric acid (GABA)ergic signaling controls the excitability of neuronal networks (Roth and Draguhn, 2012). Deficits in GABAergic neurotransmission are associated with neurodevelopmental disorders, including attention deficit / hyperactivity disorder (Edden et al., 2012), developmental co-ordination disorder (Umesawa et al., 2020), and depressive disorders (Luscher et al., 2011). GABAergic neurotransmission also plays a key role in regulating food intake (Meng et al., 2016; Xu et al., 2012). GABA is synthesized from glutamate by glutamic acid decarboxylase (GAD), an enzyme encoded by *GAD1* and *GAD2* genes coding for GAD67 and GAD65 isoforms. GAD67 produces the majority of GABA and is required for development of inhibitory neuronal circuits and maintenance of basal inhibitory firing. GAD65 synthesizes GABA to fine-tune and maintain GABAergic synapse function during neuronal activity (Baekkeskov and Kanaani, 2009; Patel et al., 2006). GAD65 is a soluble cytosolic protein, which attaches to membranes via hydrophobic modifications, with subsequent palmitoylation required for its targeting to synaptic vesicles (Baekkeskov and Kanaani, 2009). Glutamate decarboxylating activity of purified GAD65 is increased on purified synaptic vesicles (Hsu et al., 2000) and is coupled to the transport of GAD65-synthesized GABA into synaptic vesicles by the vesicular GABA transporter (VGAT) (Jin et al., 2003). Synaptic vesicles fuse with the PM to release GABA and are then reformed via the retrieval of synaptic vesicle membranes from the PM. Many peripheral proteins involved in this vesicle recycling process are recruited to synapses and then to synaptic vesicle membranes in a highly regulated manner and at different stages of the recycling process. GAD65 is co-purified with vesicles in the presence of pyridoxal 5’-phosphate, a co-factor necessary for GAD65-mediated synthesis of GABA (Jin et al., 2003), but not in its absence (Reetz et al., 1991; Takamori et al., 2006; Takamori et al., 2000), suggesting that the attachment of GAD65 to vesicles is regulated. Regulation of the synaptic targeting of GAD65 is, however, poorly understood.

In this work we show that GAD65 attaches to the PM and is removed from the PM with newly formed vesicles. NEGR1 targets GAD65 to synapses by promoting clustering of GAD65 at the presynaptic PM. NEGR1 deficiency causes GABAergic synapse loss indicating that clustering of GAD65 at the presynaptic PM is required for the maintenance of inhibitory synapses.

## Results

### NEGR1 promotes synaptic targeting of GAD65

Since NEGR1 is involved in regulating feeding behavior, we studied its distribution in cultures of hypothalamic neurons, which play a key role in regulating food intake. Confocal microscopy showed that NEGR1 was present along dendrites and axons (Fig. 1A) identified as thick tapering MAP2-positive protrusions and thin tau-enriched protrusions of a uniform diameter, respectively (Fig. 1B). The majority of synapses in these neurons (76.6 ± 2.03%, n = 20 neurons) were positive for GAD65. NEGR1 clusters co-localized with accumulations of VGAT and GAD65 indicating that NEGR1 accumulates in inhibitory synapses (Fig. 1A) and suggesting that it plays a role in regulating their formation or function.

**Figure 1.**
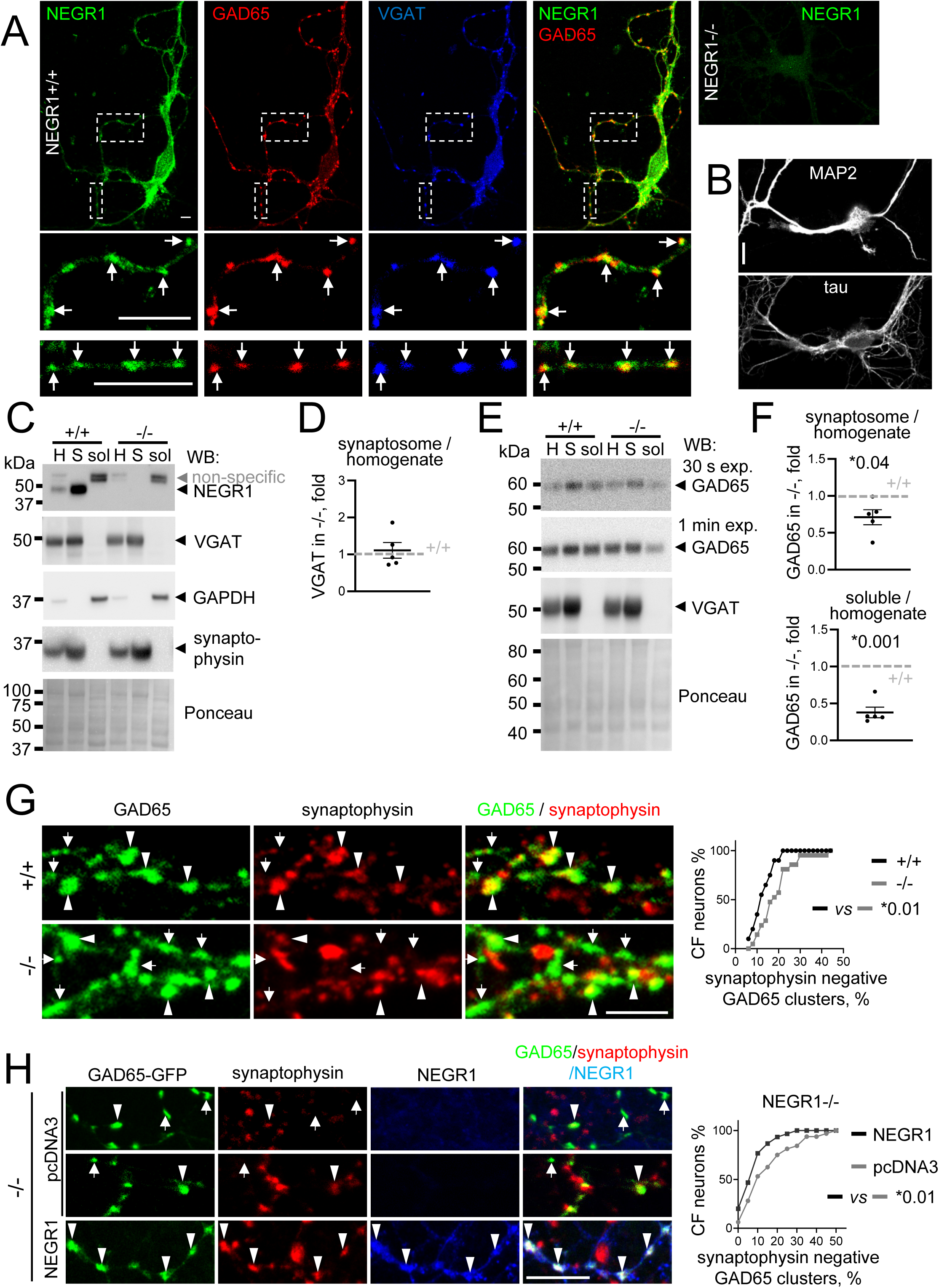
NEGR1 promotes targeting of GAD65 to synapses. **(A)** NEGR1+/+ cultured hypothalamic neuron immunolabelled for NEGR1, GAD65 and VGAT. Magnified images of the outlined areas show examples of NEGR1 clusters overlapping with GAD65 and VGAT accumulations (arrows). NEGR1-/- neurons were labelled for control. Bars, 10 µm. **(B)** NEGR1+/+ cultured hypothalamic neurons immunolabelled for a dendritic marker protein MAP2 and axonal marker protein tau. Bar, 20 µm. **(C-F)** Western blot (WB) analysis of VGAT (**C, D**) and GAD65 (**E, F**) in NEGR1+/+ and NEGR1-/- brain homogenates (H), synaptosomes (S) and soluble protein (sol) fractions. NEGR1 labelling was included in (**C**) to illustrate its enrichment in synaptosomes. Two different exposures of GAD65 labelling in (**E**) are shown to better illustrate a reduction in GAD65 levels in NEGR1-/- synaptosomes and soluble protein fractions. Labelling for GAPDH and Ponceau stain was used to control loading. Labelling for synaptophysin was used to confirm the efficiency of synaptosome isolation. Graphs show mean ± SEM changes in protein enrichment in NEGR1-/- synaptosomes and soluble protein fractions *vs* enrichment in NEGR1+/+ samples set to 1. n = 5 pairs of NEGR1+/+ and NEGR1-/- mice were analyzed. *p, one sample t test *vs* +/+. (**G**) NEGR1+/+ and NEGR1-/- cultured hypothalamic neurons immunolabeled for GAD65 and synaptophysin. (**H**) NEGR1-/- cultured hypothalamic neurons transfected with GAD65-GFP and pcDNA3 or NEGR1 immunolabelled for synaptophysin and NEGR1. In **G**, **H**, examples of synaptophysin positive (arrowheads) and negative (arrows) GAD65 clusters are shown. Bars, 5 µm. Graphs show cumulative frequency (CF) of neurons with different percentages of synaptophysin negative GAD65 clusters. *p, Mann-Whitney test (n = 20 (**G**) and 30 (**H**) neurons per group). Figure 1 source data. Contains uncropped Western blot source data for Figure 1.

Western blot analysis demonstrated that the levels of an inhibitory synapse marker protein VGAT were similar in brain homogenates of 3-month-old NEGR1+/+ and NEGR1-/- mice (Fig. 1C), while the levels of GAD65 tended to be higher in NEGR1-/- brain homogenates (Fig. 1E) (mean ± SEM fold change in NEGR1-/- *vs* NGER1+/+ was 1.17 ± 0.08 for VGAT and 1.67 ± 0.31 for GAD65 (n = 5, p = 0.06, Wilcoxon signed rank test)). NEGR1 was highly enriched in synaptosomes *vs* brain homogenates of NEGR1+/+ mice (Fig. 1C). Hence, we asked whether NEGR1 deficiency affects targeting of GAD65 to synapses. To take into account changes in total protein levels, synaptic targeting (ST) was estimated by calculating the ratio of protein levels in synaptosomes and brain homogenates in the same animal. ST_GAD65_ was reduced in NEGR1-/- *vs* NEGR1+/+ mice (Fig. 1E, F), whereas ST_VGAT_ was not changed (Fig. 1C, D). To determine whether NEGR1 promotes synaptic targeting of GAD65 by recruiting it to membranes from the soluble protein pool, the enrichment of GAD65 in soluble protein fractions relative to its total levels in brain homogenates was analyzed. Surprisingly, GAD65 enrichment in the soluble protein fraction was reduced in NEGR1-/- vs NEGR1+/+ mice (Fig. 1E, F), indicating that NEGR1 deficiency causes accumulation of GAD65 in non-synaptic membranes.

GAD65 distribution was then analyzed in cultured NEGR1+/+ and NEGR1-/- hypothalamic neurons by confocal microscopy. In NEGR1+/+ neurons, GAD65 clusters predominantly co-localized with synaptophysin accumulations (Fig. 1G), however, GAD65 clusters which did not co-localize with synaptophysin were also observed (Fig. 1G). The percentage of these non-synaptic GAD65 clusters was increased in NEGR1-/- neurons (Fig. 1G). To exclude a possibility that this effect was solely due to increased levels of GAD65 in NEGR1-/- *vs* NEGR1+/+ neurons, the synaptic targeting of GAD65- GFP was compared in NEGR1-/- hypothalamic neurons co-transfected with NEGR1 or empty pcDNA3 vector. In NEGR1-co-transfected neurons, clusters of GAD65-GFP co-localized with NEGR1 accumulations, and the percentage of synaptophysin negative GAD65-GFP clusters was reduced (Fig. 1H). Our combined observations thus indicate that NEGR1 promotes synaptic targeting of GAD65.

### Trans-interactions of NEGR1 promote synaptic targeting of GAD65

To determine whether GAD65 is targeted to synapses by trans-interactions of NEGR1, NEGR1+/+ cultured hypothalamic neurons were treated with recombinant soluble NEGR1 (sNEGR1), which binds to cell surface NEGR1 (Kim et al., 2014). Remarkably, sNEGR1 induced a strong increase in synaptic GAD65 levels (Fig. 2A). The effect was observed at 15 min after application of sNEGR1 and lasted for 24 h (the last time point analyzed) (Fig. 2A). A similar effect was found in slices of NEGR1+/+ hypothalamic tissue treated for 24 h with sNEGR1. Western blot analysis of synaptosomes from these slices showed an increase in synaptic levels of GAD65 when compared to GAD65 levels in synaptosomes from control tissues treated with bovine serum albumin (Fig. 2B). Interestingly, the overall levels of GAD65 were also increased in sNEGR1-treated slices (Fig. 2B). To analyze whether the NEGR1-dependent increase in synaptic GAD65 levels results in enhanced GABA synthesis, GABA levels in the synaptosomes were determined by mass spectrometry. Surprisingly, this analysis showed that GABA levels were reduced in synaptosomes isolated from sNEGR1-treated tissues despite the increase in GAD65 levels, indicating that NEGR1-mediated trans-interactions restrain GAD activity (Fig. 2C).

**Figure 2.**
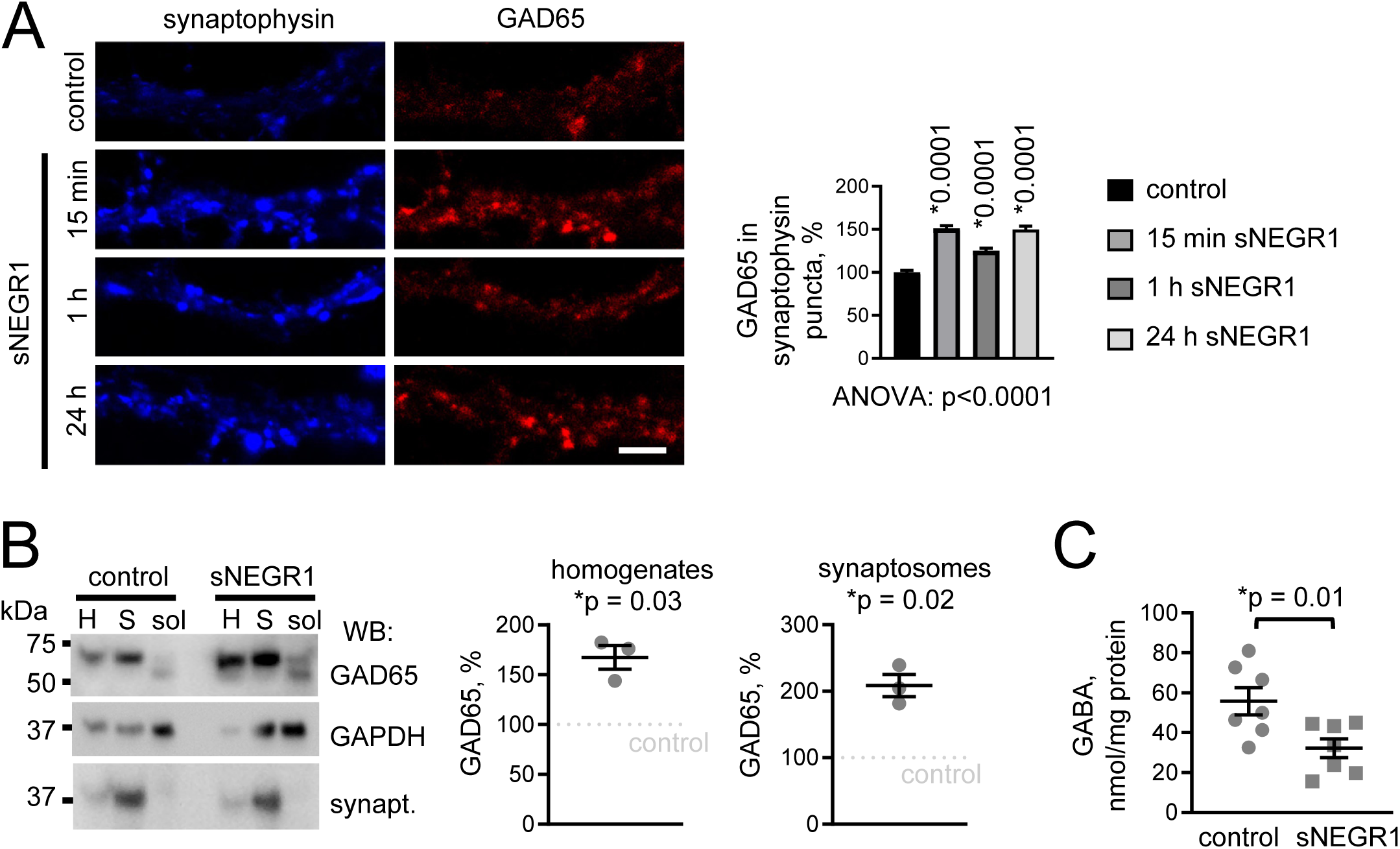
Trans-interactions of NEGR1 induce an increase in synaptic GAD65 levels. **(A)** NEGR1+/+ cultured hypothalamic neurons treated with recombinant soluble NEGR1 (sNEGR1) or mock-treated with the culture medium (control) immunolabeled for GAD65 and synaptophysin. Bar, 5 µm. Graph shows mean + SEM levels of GAD65 in synaptophysin accumulations relative to the mean value of the control set to 100% (n > 1000 boutons analyzed per group). *p, one-way ANOVA and Dunnett’s multiple comparisons test (compared to control). **(B)** Western blot (WB) analysis of GAD65 levels in homogenates (H), synaptosomes (S) and soluble protein fractions (sol) from hypothalamic tissues treated with BSA (control) or sNEGR1. GAPDH and synaptophysin served as loading controls. Graphs show mean ± SEM GAD65 levels in sNEGR1-treated tissues relative to GAD65 levels in the control set to 100% from n = 3 independent experiments. *p, one sample t test *vs* control. **(C)** Concentration of GABA in synaptosomes from hypothalamic tissues treated with BSA (control) or sNEGR1 (n = 7). *p, t test. Figure 2 source data. Contains uncropped Western blot source data for Figure 2.

### NEGR1 restrains the retrieval of synaptic vesicle membranes from the PM

GAD65 protein and mRNA levels are reduced in response to prolonged stimulation of synaptic vesicle recycling (Buddhala et al., 2012), suggesting that the regulation of GAD65 levels is coupled to synaptic vesicle recycling. Since sNEGR1 induced an overall increase in GAD65 levels, we asked whether NEGR1 regulates synaptic vesicle recycling. Activity-dependent synaptic vesicle recycling was visualized in cultured NEGR1+/+ and NEGR1-/- hypothalamic neurons by loading synaptic vesicles for 2 min with the lipophilic FM4-64 dye applied in buffer containing 47 mM K^+^, which causes PM depolarization and Ca^2+^ influx initiating synaptic vesicle fusion with the synaptic PM followed by vesicle reformation. The amount of FM4-64 taken up into synaptic boutons was increased in NEGR1-/- neurons (Fig. 3A). FM4-64 release in response to electric field stimulation was then analyzed as a measure of synaptic vesicle fusion with the PM and was found to be faster in NEGR1-/- neurons (Fig. 3A).

**Figure 3.**
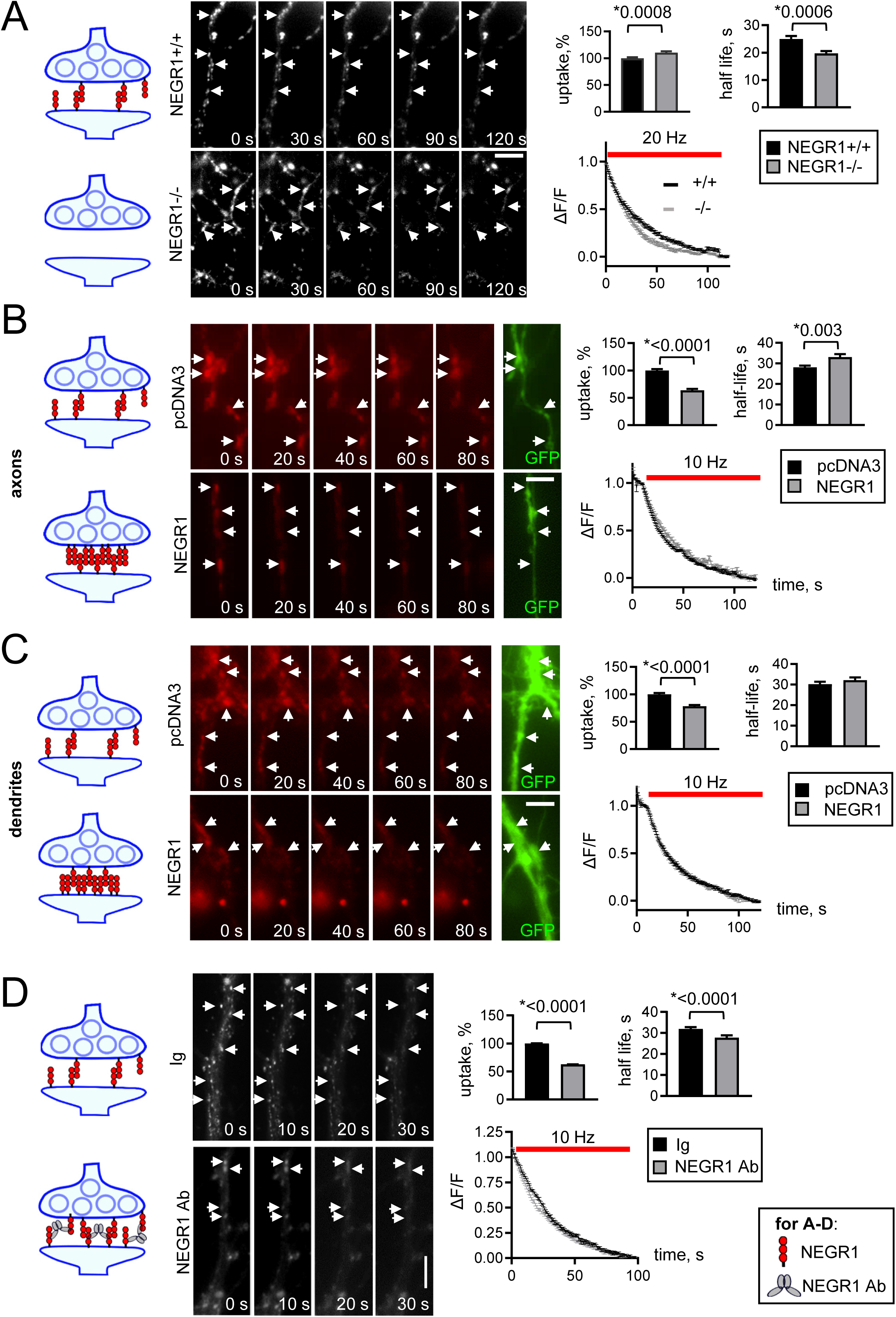
NEGR1 constrains synaptic vesicle membrane retrieval. **(A-D)** Synaptic vesicles in cultured hypothalamic neurons were loaded with FM4-64 applied for 2 min in 47 mM K+ buffer (0 s). Time lapse images show release of the dye in response to the electric field stimulation. Bar, 5 μm. Graphs show FM4-64 uptake at 0 s, changes in FM4-64 levels (ΔF/F) in synaptic boutons over time during the stimulation, and half-life of FM4-64 loss during the stimulation (mean ± SEM, *p, Mann-Whitney test). **(A)** NEGR1+/+ neurons (n = 113 boutons from 10 neurons) and NEGR1-/- neurons (n = 108 boutons from 10 neurons). **(B, C)** NEGR1+/+ neurons co-transfected with GFP and pcDNA3 or NEGR1. Axons (**B,** n = 418 boutons from 10 pcDNA3-transfected neurons, 266 boutons from 12 NEGR1-transfected neurons) and dendrites (**C**, n = 376 boutons from 12 pcDNA3-transfected neurons, 261 boutons from 13 NEGR1- transfected neurons). **(D)** NEGR1+/+ neurons treated with control non-specific immunoglobulins (Ig, n = 681 boutons from 10 neurons) or NEGR1 antibodies (NEGR1 Ab, n = 514 boutons from 10 neurons).

NEGR1 was reported to be present post-synaptically (Hashimoto et al., 2008) but was also detected in pre-synaptic compartments (Takamori et al., 2006) and was proposed to mediate trans-synaptic adhesion (Ranaivoson et al., 2019; Venkannagari et al., 2020). Hence, we investigated whether pre- or post-synaptic NEGR1 regulates synaptic vesicle recycling by determining the effects of NEGR1 overexpression in axons and dendrites of cultured NEGR1+/+ hypothalamic neurons. Control neurons were transfected with the empty pcDNA3 vector. Neurons were co-transfected with GFP, and their proximal dendrites were identified as thick tapering protrusions, while axons were identified as thin protrusions of a uniform diameter. Synapses formed by GFP-positive axons of transfected neurons on GFP-negative dendrites of non-transfected neurons were analyzed to determine the effect of presynaptic overexpression. Synapses formed by GFP-negative axons of non-transfected neurons on GFP-positive dendrites of transfected neurons were analyzed to determine the effect of postsynaptic NEGR1 overexpression. The FM4-64 uptake was reduced in synaptic boutons formed by axons of NEGR1 overexpressing *vs* control neurons (Fig. 3B) and was also reduced in synapses formed on dendrites of NEGR1 overexpressing *vs* control neurons (Fig. 3C). FM4-64 release in response to the electric field stimulation was slower in axons of NEGR1 overexpressing *vs* control neurons (Fig. 3B) and was similar in synapses formed on dendrites of NEGR1 overexpressing and control neurons (Fig. 3C).

To exclude the possibility that these changes in synaptic vesicle recycling reflect solely developmental effects of prolonged NEGR1 loss or overexpression, NEGR1+/+ cultured hypothalamic neurons were acutely treated with antibodies against NEGR1 used as a specific NEGR1 ligand, a method used to cluster and mimic trans-interactions of cell adhesion molecules at the cell surface (Sheng et al., 2015; Sheng et al., 2019). Control neurons were incubated with non-specific immunoglobulins (Ig). The FM4-64 uptake was strongly reduced in NEGR1 antibody- *vs* Ig-treated neurons (Fig. 3D). Unexpectedly, FM4-64 release in response to the electrical field stimulation was slightly faster in NEGR1 antibody-treated neurons (Fig. 3D).

Altogether, our data indicate that both pre- and post-synaptic NEGR1 restrain the retrieval of synaptic vesicle membranes from the synaptic PM, while only presynaptic NEGR1 also plays a role in regulating fusion of synaptic vesicles with the synaptic PM.

### Inhibition of synaptic vesicle membrane retrieval promotes GAD65 clustering at the synaptic PM

To determine whether a restraint of synaptic vesicle membrane retrieval influences the synaptic localization of GAD65, the association of GAD65 clusters with synaptophysin accumulations was analyzed in NEGR1+/+ cultured hypothalamic neurons treated for 15 min with dynasore. This inhibitor of dynamin selectively suppresses synaptic vesicle membrane retrieval after action potential-induced neurotransmitter release but does not affect spontaneous synaptic vesicle trafficking (Chung et al., 2010). Control neurons were treated with vehicle (DMSO). The numbers of synaptophysin-negative GAD65 clusters were reduced in dynasore-treated *vs* control neurons (Fig. 4A). sNEGR1 induced a similar effect in a non-additive manner with dynasore (Fig. 4A). Interestingly, tetanus toxin, which blocks fusion of synaptic vesicles with the PM, also reduced the percentage of synaptophysin-negative GAD65 clusters in a non-additive manner with sNEGR1 (Fig. 4B).

**Figure 4.**
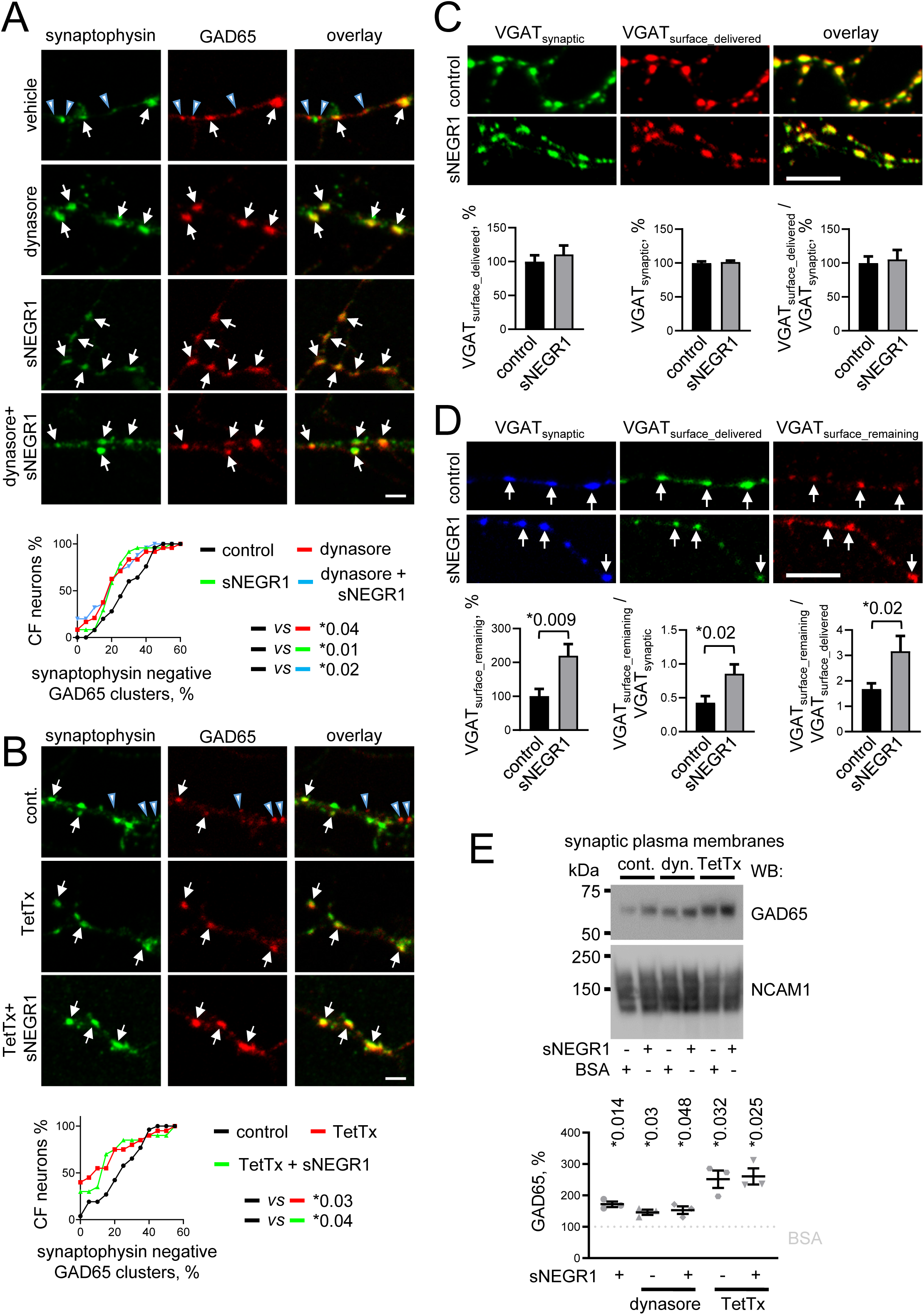
Trans-interactions of NEGR1 promote clustering of GAD65 at the synaptic PM by constraining the retrieval of synaptic vesicle membranes. **(A, B)** NEGR1+/+ cultured hypothalamic neurons treated with sNEGR1 and dynasore (**A**) or TetTx (**B**) as indicated. Control neurons were mock treated with vehicle (DMSO in **A**, water in **B**). Neurons were immunolabeled for GAD65 and synaptophysin. Examples of synaptophysin positive (arrows) and negative (arrowheads) GAD65 clusters are shown. Bars, 5 µm. Graphs show cumulative frequency (CF) of neurons with increasing percentages of synaptophysin negative GAD65 clusters. *p, Mann-Whitney test (n = 20-26 neurons per group). **(C, D)** sNEGR1-treated and control culture medium-treated NEGR1+/+ cultured hypothalamic neurons incubated live with antibodies against the lumenal domain of VGAT. The antibodies were detected with secondary antibodies applied before (VGAT_surface_remaining_) and after (VGAT_surface_delivered_) permeabilization of membranes with detergent. Neurons were co-labelled with antibodies against the cytoplasmic domain of VGAT (VGAT_synaptic_). Bar, 5 μm. Graphs show mean + SEM labelling intensities of VGAT_synaptic_ and VGAT_surface_delivered_ (**C**), VGAT_surface_remaining_ (**D**), and their ratios (n = 40 neurons in **C**, 20 neurons in **D**). Control mean intensity was set to 100%. **(E)** Western blot analysis of GAD65 levels in synaptic PM from hypothalamic tissues treated with BSA or sNEGR1 applied either alone (cont.) or together with dynasore (dyn.) or TetTx. NCAM1 served as loading control. Graph shows mean ± SEM GAD65 levels relative to the levels in BSA only treated tissues set to 100% from n = 3 independent experiments. Samples from each experiment were analyzed by Western blot twice and the data were averaged. *p, one sample t test *vs* BSA only treated tissues. Figure 4 source data. Contains uncropped Western blot source data for Figure 4.

Synaptic vesicle membrane retrieval is coupled to vesicle fusion with the PM (Haucke et al., 2011). To determine whether sNEGR1 inhibits fusion of synaptic vesicles with the PM or retrieval of their membranes from the PM in GABAergic synapses, live cultured hypothalamic neurons pretreated with sNEGR1 or culture medium for control were incubated for 30 min with antibodies against the lumenal domain of VGAT. The antibodies bind to the lumenal domain of VGAT when it is exposed at the cell surface after fusion of synaptic vesicles with the PM. The antibodies then either remain at the cell surface or are taken up into vesicles reformed via membrane retrieval from the PM. The pool of antibodies remaining at the cell surface (VGAT_surface_remaining_), which is inversely proportional to the retrieval rate, was visualized with secondary antibodies applied to neurons which were not permeabilized with detergent. The total pool of antibodies at the cell surface and in newly formed vesicles (VGAT_surface_delivered_), which is proportional to the rate of synaptic vesicle fusion with the PM, was visualized with secondary antibodies applied to fixed and detergent-permeabilized neurons (Fig. 4C, D). The total synaptic pool of VGAT (VGAT_synaptic_) was visualized by co-labeling detergent-permeabilized neurons with antibodies against the cytoplasmic domain of VGAT. VGAT_surface_delivered_ and VGAT_synaptic_ were similar in sNEGR1-treated and control neurons (Fig. 4C), whereas VGAT_surface_remaining_ was increased in sNEGR1-treated neurons (Fig. 4D), indicating that sNEGR1 did not affect the fusion of vesicles with the PM but inhibited the retrieval of synaptic vesicle membranes from the PM.

Inhibition of synaptic vesicle membrane retrieval causes clustering of synaptic vesicle proteins in the presynaptic PM (Dason et al., 2014). To test whether sNEGR1 induces accumulation of GAD65 in the presynaptic PM, synaptosomes isolated from slices of hypothalamic tissue were used to obtain synaptic PMs isolated after synaptosome lysis with osmotic shock, which releases synaptosomal contents including synaptic vesicles. Western blot analysis demonstrated that GAD65 was co-isolated with the synaptic PMs (Fig. 4E). In slices treated with sNEGR1 for 24 h, the levels of GAD65 co-isolated with the synaptic PMs were ∼2-fold higher than in membranes from slices treated with bovine serum albumin for control (Fig. 4E). Dynasore induced a similar increase in GAD65 levels in the synaptic PMs in a non-additive manner with sNEGR1 (Fig. 4E). An increase in GAD65 levels co-isolated with the synaptic PMs was also found in slices treated with tetanus toxin (Fig. 4E), indicating that fusion of vesicles with the PM is not required for clustering of GAD65 at the synaptic PM and that GAD65 is not delivered to the synaptic PM by synaptic vesicles.

### GAD65 attaches to the PM and NEGR1 promotes its recruitment to the PM-derived vesicles in CHO cells

Our finding that GAD65 is co-isolated with the synaptic PM contrasts with extensive microscopy data in different cell types showing that GAD65 does not accumulate at the PM. It is, however, inherently difficult to visualize transient interactions of peripheral proteins such as GAD65 with the PM in intact cells. We attempted to achieve this by using bimolecular fluorescence complementation (BiFC) to simultaneously capture and visualize GAD65 in close proximity to the PM via irreversible reconstitution of the Venus fluorescent protein. As a membrane localized sensor, we engineered an LCK-VN protein consisting of the N-terminal fragment of Venus fused to the first 26 amino acids of human LCK containing sites for myristoylation and palmitoylation which target it to the PM (Fig. 5A). The sensor was used to detect the PM attachment of GAD65 fused to the complementary C-terminal fragment of Venus (GAD65-VC). Reconstitution of Venus from the VC and VN fragments brought in close proximity to each other by the attachment of GAD65 to the PM captures GAD65 at the site of PM attachment and simultaneously visualizes it by producing fluorescence (Fig. 5A). The crowded environment of synaptic boutons does not allow unambiguous determination of whether proteins are bound to the PM rather than synaptic vesicles. Therefore, we used cultured CHO cells as a model system to visualize GAD65 interactions with the PM in vesicle free PM areas. Labelling of LCK-VN transfected CHO cells with antibodies against GFP, which detect the VN fragment, showed that this sensor was localized at the PM and intracellular accumulations partially overlapping with the Golgi marker GM130 (Fig. 5B). Co-transfection of cells with LCK-VN and non-mutated GAD65WT-VC produced BiFC fluorescence, which accumulated in the perinuclear region (Fig. 5C) where GAD65 tightly attaches to the Golgi (Kanaani et al., 2008). In addition, clusters of BiFC fluorescence were found at the PM and in small accumulations in the cytosol (Fig. 5C). A closer analysis of cross-sections of the 3D reconstructed transfected cells showed that BiFC signals at the PM were found in small clusters or vesicle-like structures (Fig. 5D). A similar level of BiFC fluorescence was observed in CHO cells transfected with GAD65(C30,45A)-VC mutant (Fig. 5C), in which cysteines responsible for GAD65 palmitoylation (Kanaani et al., 2008) were mutated to alanine. The BiFC signal produced by the GAD65(C30,45A)-VC mutant was also found at the PM (Fig. 5C), indicating that palmitoylation is not required for the attachment of GAD65 to the PM. The BiFC signal was dramatically reduced in cells co-transfected with GAD65(24-31A)-VC (Fig. 5C), in which amino acids 24-31 responsible for binding to membranes (Shi et al., 1994) were substituted for alanine. In cells co-transfected with NEGR1, cell surface clusters of NEGR1 were found at sites of BiFC fluorescence accumulations (Fig. 5E). A subpopulation of PM-localized BiFC fluorescence clusters co-localized with clathrin accumulations suggesting that the attachment of GAD65 to the PM triggers its recruitment to the clathrin-coated pits (Fig. 5F). To determine whether NEGR1 plays a role in recruiting GAD65 to the vesicles derived from the PM, CHO cells were incubated live with FM4-64 to load the dye in and visualize newly formed vesicles. Since stabilization of the PM attachment via BiFC may influence endocytosis, GAD65-GFP was used in these experiments to visualize GAD65 on the vesicles. The levels of FM4-64 uptake were reduced, however, the percentages of FM4-64-loaded vesicles co-localizing with GAD65 accumulations were increased in NEGR1-transfected vs control pcDNA3-transfected cells (Fig. 5G). Altogether, these observations indicate that GAD65 attaches to the PM and NEGR1 promotes its targeting to the PM-derived vesicles. Our data also further suggests that this sorting requires a restraint of the retrieval of membranes from the cell surface.

**Figure 5.**
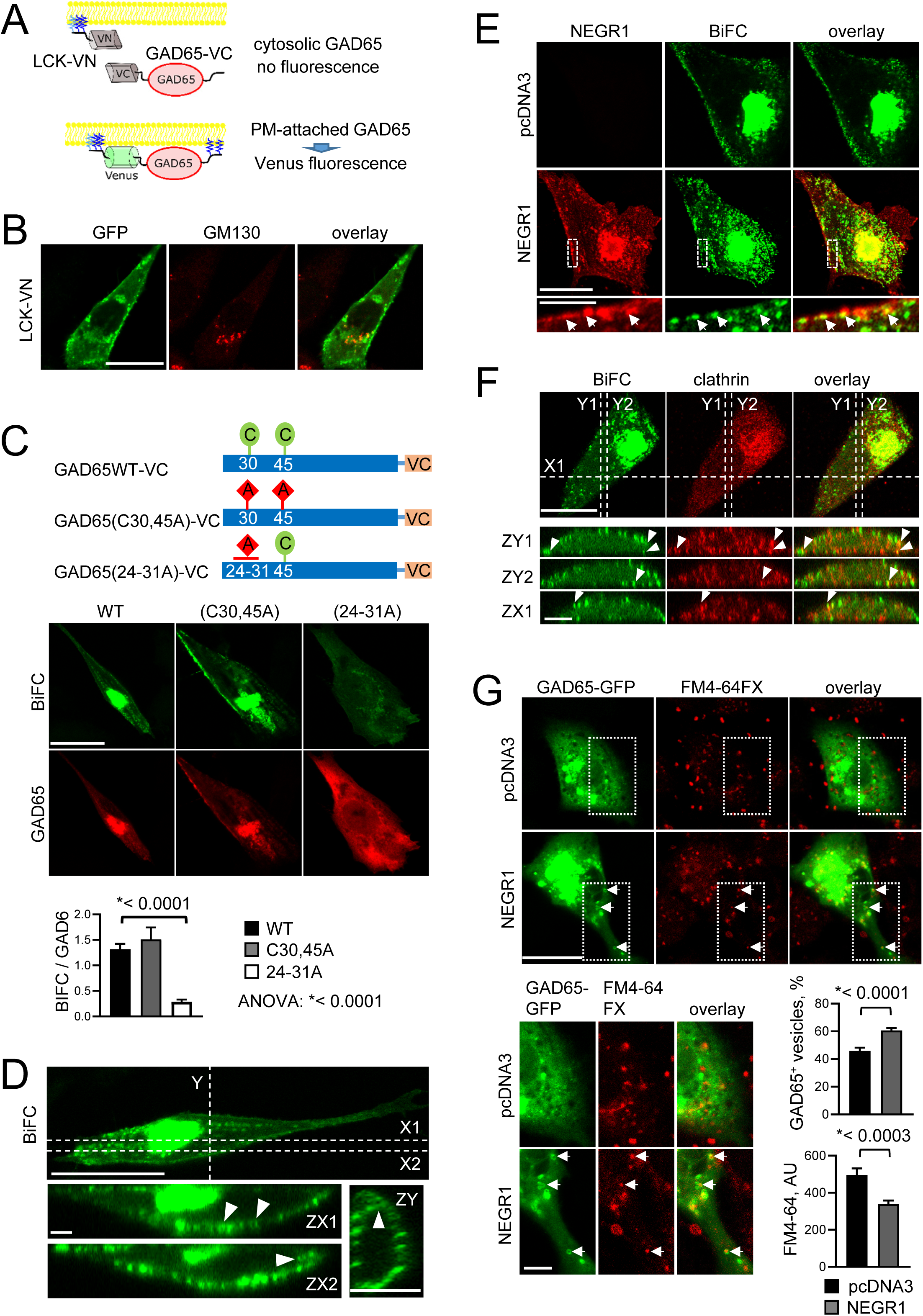
NEGR1 promotes the association of GAD65 with vesicles formed at the PM. **(A)** Diagram illustrating the principle of detection of the PM attachment of GAD65 using BiFC. **(B)** LCK-VN-transfected CHO cell labelled with VN-recognizing GFP antibodies and co-labelled for GM130. **(C)** BiFC assay with CHO cells co-transfected with LCK-VN and GAD65-VC proteins shown on the diagram. Immunolabelling for GAD65 was used for normalization. Graph shows mean ± SEM BiFC / GAD65 ratio. *p, one-way ANOVA and Dunnett’s multiple comparisons test (n = 15 cells per group). **(D)** BiFC fluorescence distribution in the XY confocal slice and ZX and ZY sections along the dashed lines through the 3D reconstructed GAD65WT-VC-co-transfected cell shown in **C**. Note PM localization of the BiFC signals in ZX and ZY sections. Arrowheads show vesicle-like structures at the PM. **(E)** CHO cells co-transfected with LCK-VN, GAD65WT-VC and either pcDNA3 or NEGR1. Cells were co-labelled for NEGR1. Arrows show clusters of NEGR1 co-localized with BiFC signals. **(F)** CHO cell co-transfected with LCK-VN and GAD65WT-VC co-labelled for clathrin. Arrowheads in ZX and ZY sections through the 3D reconstructed cell show BiFC signals co-localized with clathrin accumulations at the PM. **(G)** FM4-64 loaded vesicles in CHO cells co-transfected with GAD65-GFP and either pcDNA3 or NEGR1. High magnification images show areas outlined with dashed boxes. Arrows show examples of FM-loaded vesicles co-localized with GAD65-GFP. Graph shows mean ± SEM percentages of FM-loaded vesicles co-localized with GAD65-GFP and FM4-64 levels. *p, Mann-Whitney test (n = 42 cells per group). Bars, 20 µm (low magnification), 5 µm (high magnification) **(B-G)**.

### NEGR1 promotes the assembly of lipid rafts at the PM and in synapses

Disruption of lipid rafts blocks synaptic clustering of GAD65 resulting in formation of non-synaptic GAD65 clusters (Kanaani et al., 2002) resembling those found in NEGR1-/- neurons (Fig. 1G). NEGR1 associates with lipid rafts via its GPI-anchor and is involved in cholesterol transport (Kim et al., 2017). We therefore asked whether NEGR1 regulates synaptic clustering of lipid microdomains. The targeting of cholesterol and ganglioside GM4 to the PM was increased in NEGR1-transfected *vs* control pcDNA3 vector-transfected CHO cells (Fig. 6A, B) indicating that NEGR1 promotes the assembly of lipid rafts at the PM. A dot blot analysis showed that the enrichment of lipid raft-localized gangliosides and phosphatidylinositol 4,5-bisphosphate in synaptosomes *vs* homogenates was strongly reduced in NEGR1-/- *vs* NEGR1+/+ mice (Fig. 6C), indicating that NEGR1 can target GAD65 to synapses by inducing synaptic clustering of lipid rafts, which can also restrain the retrieval of synaptic vesicle membranes since synaptic vesicle proteins synaptophysin and VAMP2 bind to the PM localized cholesterol (Thiele et al., 2000).

**Figure 6.**
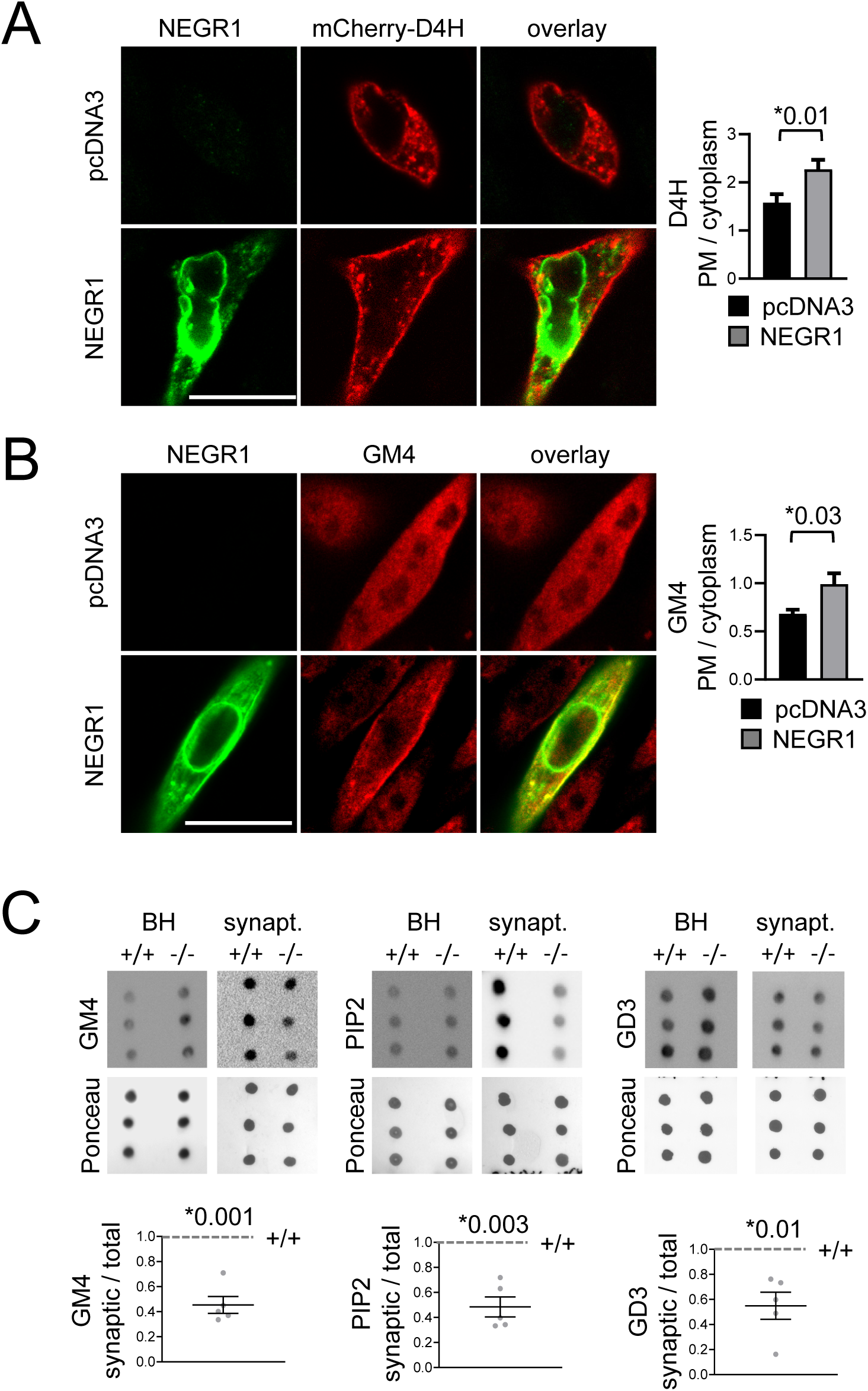
NEGR1 promotes the assembly of lipid rafts at the PM and in synapses. **(A, B)** CHO cells transfected with pcDNA3 or NEGR1 were either co-transfected with mCherry-D4H to visualize cholesterol **(A)** or immunolabelled for ganglioside GM4 **(B)**. Bars, 20 µm. Graphs show ratios of D4H and GM4 labelling levels at the PM and in cytoplasm (mean + SEM, n = 20). *p, Mann Whitney test. **(C)** Lipid raft markers gangliosides GM4 and GD3 and phosphatidylinositol 4,5-bisphosphate (PIP2) analyzed in triplicate by dot blot in brain homogenates (BH) and synaptosomes (synapt.) from a representative pair of NEGR1+/+ and NEGR1-/- littermates. Graphs show mean + SEM synaptic enrichments of the lipids in NEGR1-/- mice relative to the levels in NEGR1+/+ mice set to 1. n = 5 pairs of NEGR1+/+ and NEGR1-/- mice were analyzed. *p, one sample t test *vs* +/+. Figure 6 source data. Contains dot blot source data for Figure 6.

### The association of GAD65 with vesicles promotes GABA synthesis

The trafficking of GAD65 to post-Golgi vesicular membranes is controlled by its palmitoylation cycle (Kanaani et al., 2008). We assessed whether a palmitoylation-deficient mutant of GAD65, which is capable of firmly anchoring to ER/cis-Golgi but incapable of anterograde trafficking from cis-Golgi to TGN and post-Golgi vesicles (Phelps et al., 2016), would exhibit altered enzyme activity. Pancreatic beta cells synthesize and secrete large quantities of GABA, making them excellent candidates for studying GAD enzyme activity (Menegaz et al., 2019). INS-1 and MIN6 beta cell lines have lost the expression of endogenous GAD65 and GAD67 (Cianciaruso et al., 2017). We thus used INS-1 beta cells to study the GAD enzyme activity of cells transfected with GAD65-GFP relative to the palmitoylation deficient mutant GAD65(C30,45A)-GFP in the absence of a background of endogenous GABA biosynthesis. As previously observed (Kanaani et al., 2008; Phelps et al., 2016), GAD65-GFP distributed between cytosol, ER, Golgi, and post-Golgi vesicles, while GAD65(C30,45A)-GFP was distributed only to the cytosol, ER, and cis-Golgi (Fig. 7A). HPLC analysis of amino acids from INS-1E cell lysate showed a 50% decrease in the cellular GABA content for GAD65(C30,45A)-GFP and a significant increase in glutamate, the precursor to GABA synthesis (Fig. 7B-D). These data demonstrate that palmitoylation-dependent vesicular trafficking of GAD65 is correlated with higher levels of GABA biosynthesis.

**Figure 7.**
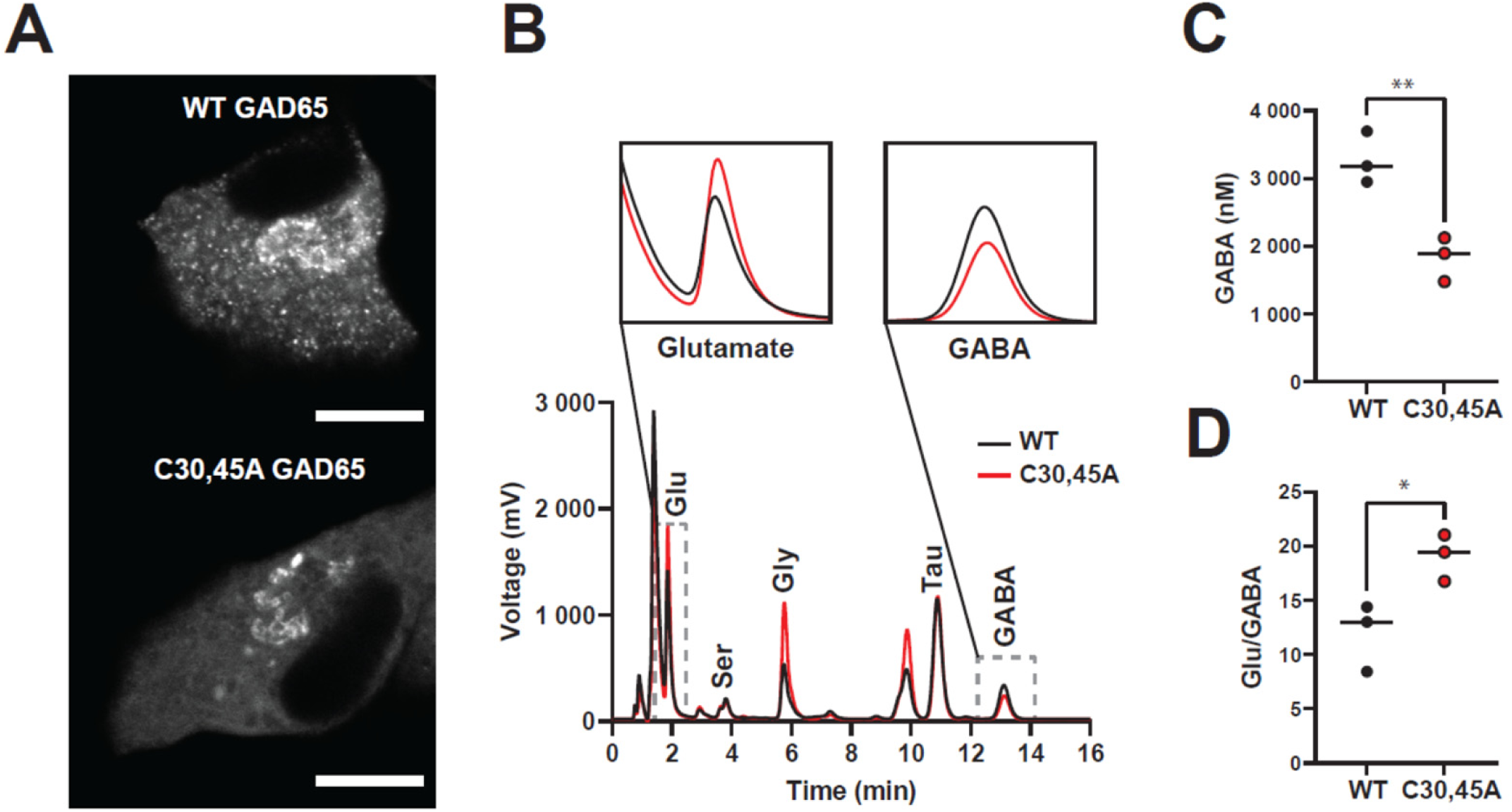
Vesicular localization of GAD65 promotes GAD65-mediated GABA synthesis. **(A)** Representative images of wild-type and palmitoylation mutant GAD65 expressed in INS-1E cells. Bar, 10 μm. **(B)** Chromatogram and isolated peaks showing amino acid content of wild-type vs. mutant. **(C)** GABA content of transfected INS-1E cells. *p = 0.008, t test (n = 3). **(D)** Glutamate to GABA ratios in transfected INS-1E cells. *p = 0.03, t test (n = 3).

### The density of GABAergic synapses is reduced in 7-8-month-old NEGR1-/- mice

In cultures of disassociated hypothalamic neurons, NEGR1 was particularly strongly expressed in axons and synaptic boutons of neuropeptide Y (NPY) / GAD65 positive neurons when compared to other neurons present in these cultures (Fig. 8A). NPY positive neurons and their projections are present at high density in the ARC (Suyama and Yada, 2018). Analysis of NEGR1+/+ and NEGR1-/- brain sections by confocal microscopy showed that the density of VGAT/GAD65 positive inhibitory synapses was strongly reduced in the ARC of NEGR1-/- *vs* NEGR1+/+ mice (Fig. 8B). In NEGR1-/- mice, the density of VGAT/GAD65 positive synapses was also reduced in the CA3 region of the hippocampus (Fig. 8C), which also contains NPY-positive interneurons (Kuruba et al., 2011). Altogether, our data indicate that NEGR1 deficiency causes a loss of inhibitory synapses.

**Figure 8.**
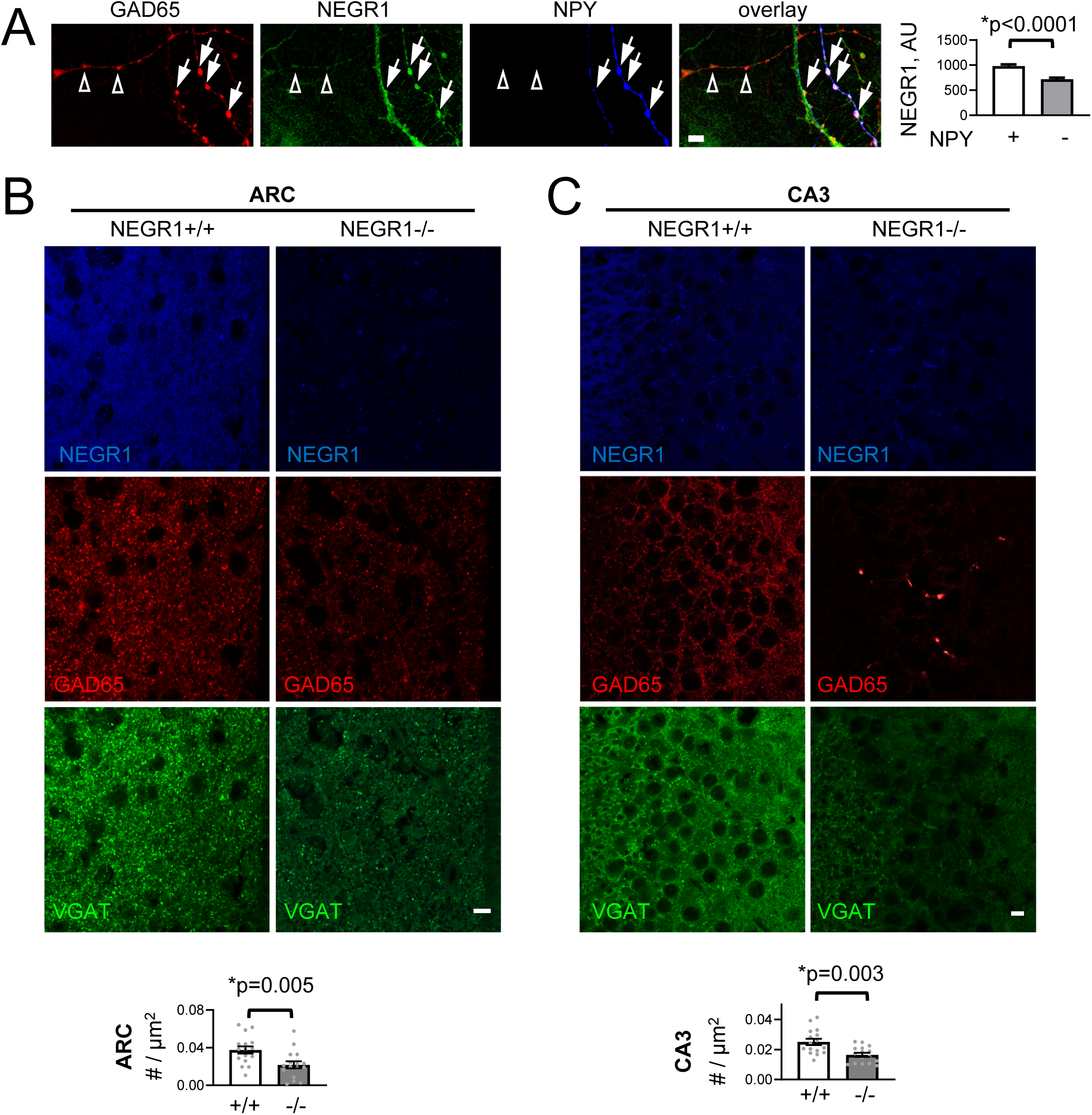
NEGR1 deficiency causes loss of inhibitory synapses. (**A**) Neurites of NEGR1+/+ cultured hypothalamic neurons immunolabelled for NEGR1, GAD65 and NPY. Arrows show examples of NEGR1 clusters overlapping with NPY positive GAD65 accumulations. Open arrowheads show NPY negative GAD65 accumulations with lower levels of NEGR1. Bar, 5 µm. Graph shows mean ± SEM levels of NEGR1 in NPY positive (+) and negative (-) GAD65 accumulations (n > 100). AU – arbitrary units defined as pixel values of 16-bit gray scale images. *p, unpaired t test. (**B, C**) Confocal images of the ARC (**B**) and pyramidal cell layer in the hippocampal CA3 region (**C**) in brain sections of NEGR1+/+ and NEGR1-/- littermates immunolabelled for NEGR1, GAD65 and VGAT. Bar, 10 µm. Graphs show mean ± SEM densities of VGAT accumulations. *p, Mann-Whitney test.

### NEGR1-/- mice show reduced motivation for a highly palatable food reward

GABAergic neurotransmission is involved in regulation of the motivational aspects of food reinforced behaviors and is sensitized by intake of palatable food (Newman et al., 2013). To determine whether NEGR1 deficiency affects motivation for food, 7-8-month-old NEGR1+/+, NEGR1-/- and NEGR1+/- mice were tested using an instrumental conditioning task. The mice were initially allowed to eat sucrose pellets released into a magazine at a rate of 1 pellet / min over 30 min. Analysis of these initial magazine training sessions performed over 6 days showed that the mice of all genotypes initially consumed similar numbers of pellets per session (Fig. 9A). Over time, the numbers of pellets consumed per session increased for NEGR1-/- but not NEGR1+/+ mice with an intermediate effect observed for NEGR1+/- mice (Fig. 9A). The mice were then trained to press a lever to obtain a sucrose pellet. In this fixed ratio task, mice of all genotypes pressed the lever with approximately the same frequency and consumed similar numbers of pellets per session (Fig. 9B). Finally, the mice were tested using a progressive ratio schedule of reinforcement, where the number of lever presses required to obtain the pellet progressively increased across the session. When the number of lever presses required to obtain a single sucrose pellet exceeded motivation for the pellet, the mice stopped responding and this was considered the break point. This analysis showed that breakpoint values were similar in all genotypes on the first day of tests (Fig. 9C), and progressively increased in NEGR1+/+ mice over three days of tests (Fig. 9C, D). Accordingly, the number of pellets delivered per session and the number of active lever presses per session also increased in NEGR1+/+ group (Fig. 9E). This sensitization was inhibited in NEGR1+/- and NEGR1-/- mice (Fig. 9C-E). Hence, NEGR1 deficiency does not negatively impact on the palatability of the food pellet or learning around the instrumental task, but rather specifically affects motivation for highly palatable food rewards.

**Figure 9.**
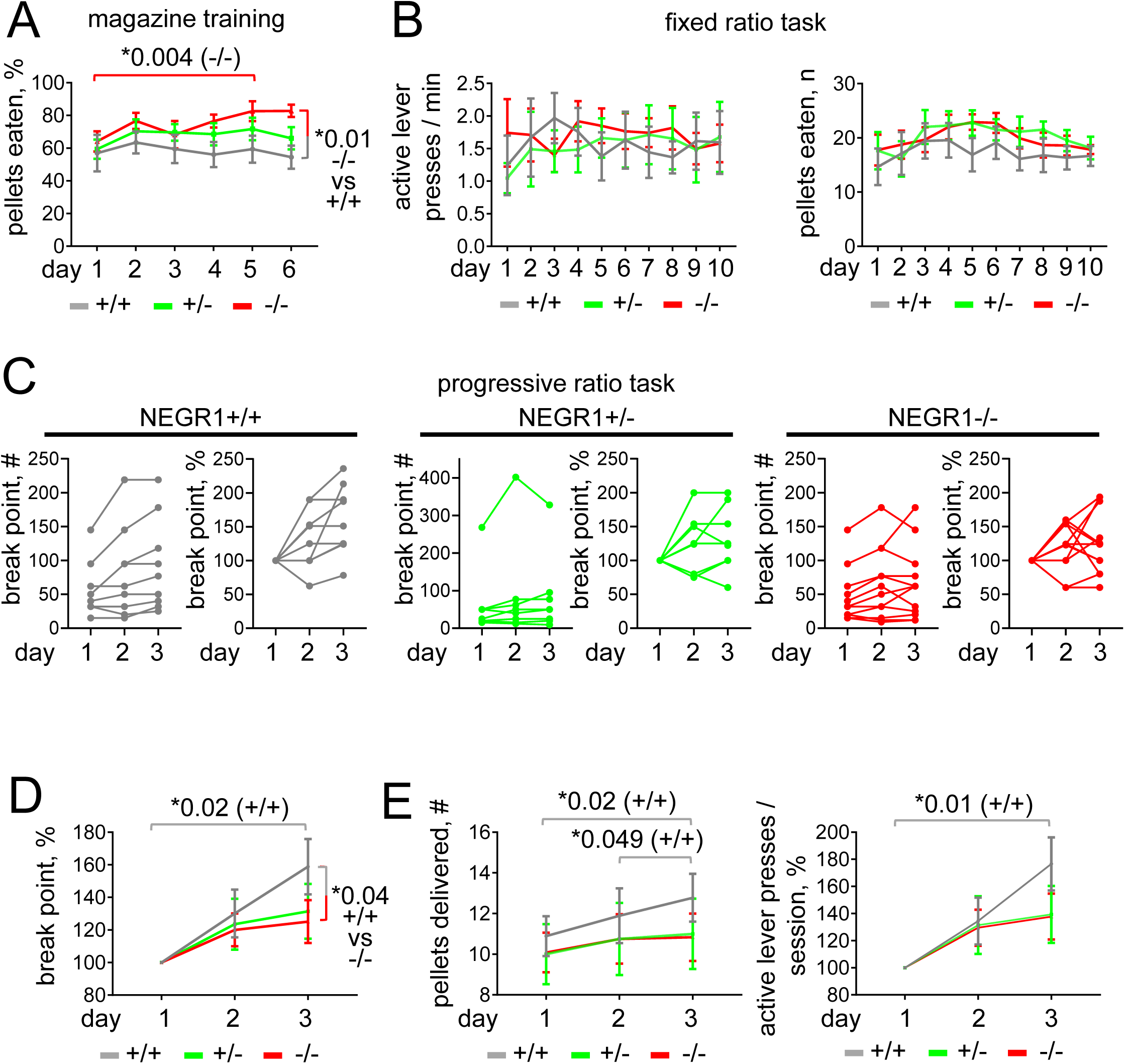
NEGR1 deficiency reduces sensitization of the reinforcing effects of food. (**A**) Mean ± SEM percentages of pellets eaten over 30 min during magazine training sessions. One pellet was released into the magazine each minute. (**B**) Mean ± SEM numbers of active lever presses and pellets eaten during the fixed ratio task. Each active lever press led to the release of a pellet. (**C**) Individual progressive ratio break point values measured over three consecutive days in the progressive ratio task. Lines connect data for individual mice. Graphs showing data normalized to the values on day 1 set to 100% are included to illustrate an increase in the progressive ratio break point in NEGR1+/+ mice. (**D**) Mean ± SEM of the normalized values shown in **C**. (**E**) Mean ± SEM numbers of pellets delivered and active lever presses in the progressive ratio task. The numbers of active lever presses were normalized to the number on day 1 set to 100%. In **A**-**E**, n = 9 NEGR1+/+, 8 NEGR1+/- and 12 NEGR1-/- mice were analyzed. *p, repeated measures two-way ANOVA and Tukey’s multiple comparisons test.

### A high fat diet causes an increase in NEGR1 levels affecting subsynaptic localization of GAD65 and synapse maintenance

A high fat diet (HFD) causes a reduction in GABA levels at least in some brain regions (Hassan et al., 2018; Sandoval-Salazar et al., 2016). Hence, we asked whether levels of NEGR1 are influenced by HFD. Western blot analysis demonstrated that levels of NEGR1 were increased in brain homogenates and synaptosomes of HFD- *vs* chow-fed mice (Fig. 10A, B). NEGR1 can be shed by ADAM10, which releases its ∼50 kDa cleavage product (Pischedda and Piccoli, 2015). The levels of this product in the soluble protein fraction were, however, below the detection limit indicating low levels of the ADAM10-mediated cleavage of NEGR1 in the mature brain (Fig. 10A). Prolonged exposure revealed a soluble fragment of NEGR1 at ∼27 kDa most likely representing the first and second Ig domains of NEGR1 (Fig. 10D, predicted molecular weight ∼24 kDa), since we used antibodies recognizing aa170-200 in the second Ig domain of NEGR1 (sc137625, Santa Cruz Biotechnology). The levels of this fragment were also increased in brains of HFD-fed mice (Fig. 10A, C) indicating that an increase in NEGR1 levels was not caused by reduced proteolysis. An increase in NEGR1 levels in HFD-fed mice was accompanied by an ∼50% increase in GAD65 levels co-purified with the synaptic PM (Fig. 10E) and ∼2-fold decrease in the concentration of GABA in synaptosomes (Fig. 10F) resembling effects in sNEGR1-treated tissues (Fig. 2C, 4E). The levels of GAD65 were reduced in brain homogenates and synaptosomes of HFD- *vs* chow-fed mice (Fig. 10G), possibly reflecting synapse loss caused by the HFD (Bocarsly et al., 2015). Selective loss of inhibitory but not excitatory synapse markers is found in dynamin 1 and 3 double knock-out neurons (Raimondi et al., 2011), indicating that defects in the retrieval of synaptic vesicle membranes affect the maintenance of inhibitory synapses. To determine whether the NEGR1 overexpression-induced decrease in synaptic vesicle membrane retrieval (Fig. 3B, C) causes changes in synapse numbers, synaptophysin levels were measured along dendrites and axons of NEGR1- and control pcDNA3-transfected cultured hypothalamic neurons. Synaptophysin levels were strongly reduced along axons and dendrites of NEGR1-overexpressing neurons (Fig. 10H). Altogether, our data indicate that NEGR1 is overexpressed in brains of HFD-fed mice, and an increase in levels of this protein correlates with reduced GABA synthesis and can cause a loss of inhibitory synapses.

**Figure 10.**
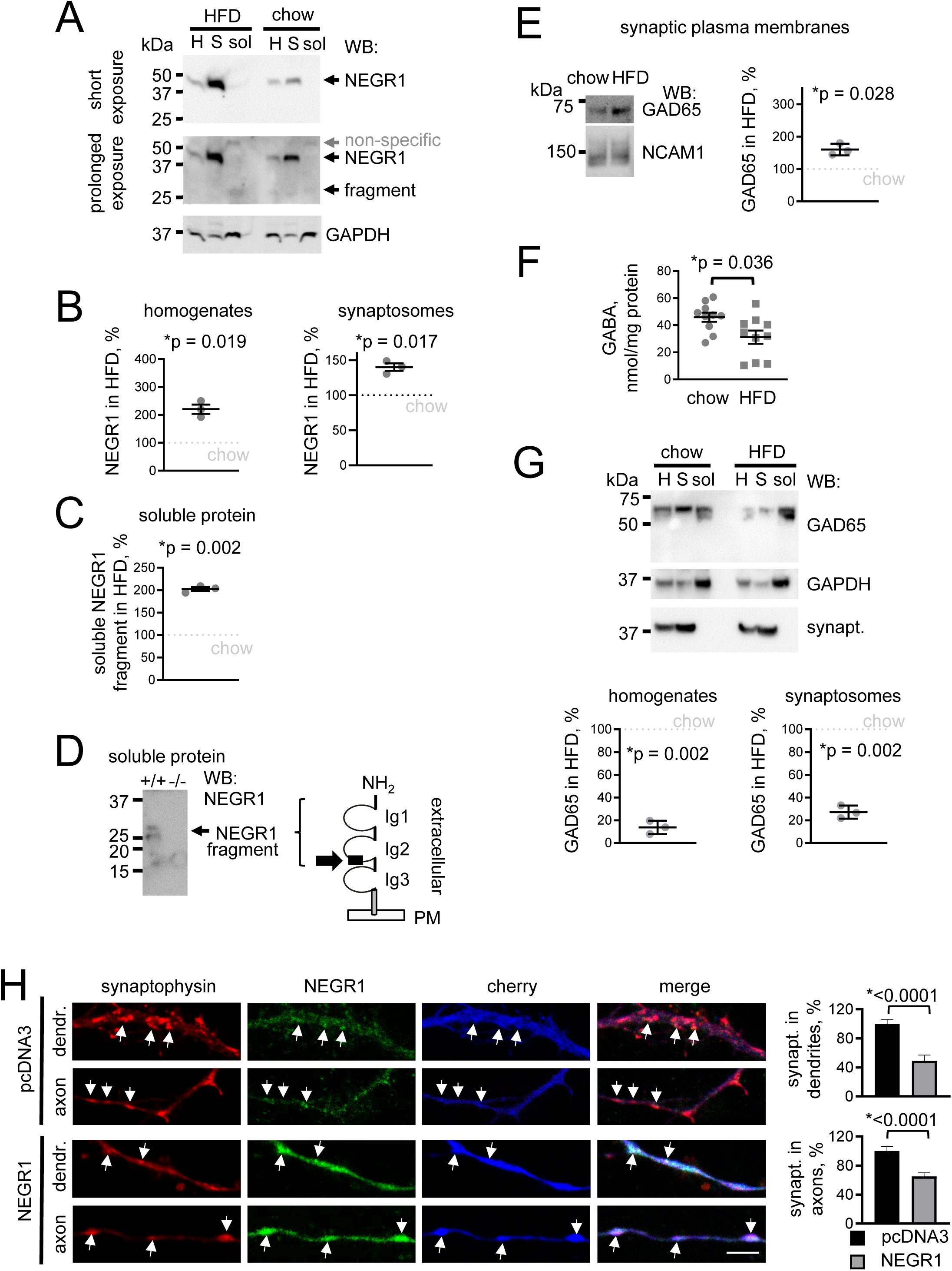
Increased NEGR1 levels correlate with a reduction in synaptic GABA and GAD65 levels and GAD65 accumulation at the synaptic PM in brains of mice fed a high fat diet. **(A-C)** Western blot (WB) analysis of NEGR1 levels in homogenates (H), synaptosomes (S) and soluble protein fractions (sol) from brains of chow and high fat diet (HFD) fed mice. GAPDH served as loading control. Graphs show levels of full length NEGR1 in brain homogenates and synaptosomes (**B**) and soluble NEGR1 fragments in the soluble protein fraction (**C**) from HFD fed mice relative to the levels in chow diet fed mice set to 100%. n = 3 mice per group. (**D**) WB analysis of the soluble protein fractions from NEGR1+/+ and NEGR1-/- brains with NEGR1 antibodies. Note that the NEGR1 fragment is detected only in the NEGR1+/+ lane. The diagram shows the structure of NEGR1 consisting of three Ig-domains attached to the PM via a GPI anchor. The site recognized by the antibody used for Western blot analysis is marked with a square. (**E**) WB analysis of GAD65 levels in synaptic PM from chow and HFD fed mice. NCAM1 served as loading control. Graph shows levels in HFD membranes relative to chow control set to 100%. n = 3 mice per group. **(F)** Concentration of GABA in synaptosomes from the brains of chow and HFD fed mice (n = 10). **(G)** WB analysis of GAD65 levels in homogenates (H), synaptosomes (S) and soluble protein fractions (sol) from brains of chow and HFD fed mice. GAPDH and synaptophysin served as loading controls. Graphs show levels of GAD65 in brain homogenates and synaptosomes of HFD fed mice relative to the levels in chow fed mice set to 100%. n = 3 mice per group. **(H)** Axons and dendrites of NEGR1+/+ cultured hypothalamic neurons co-transfected with cherry and control pcDNA3 vector or NEGR1. Neurons were immunolabelled for synaptophysin and NEGR1. Note reduced numbers of synaptophysin puncta (arrows) along the axons and dendrites of NEGR1-overexpressing neurons. Bar, 5 µm. Graphs show synaptophysin labelling intensities along dendrites (n = 14 - 24) and axons (n = 31 - 43). Mean of control was set to 100%. In **B, C, E-H**, mean ± SEM values are shown. *p, one sample t test *vs* chow (**B, C, E, G**) or Mann Whitney test (**F, H**). Figure 10 source data. Contains uncropped Western blot source data for Figure 10.

## Discussion

We report that NEGR1 promotes the synaptic targeting of GAD65, a GABA synthesizing enzyme. We demonstrate that GAD65 attaches to the synaptic PM and show that trans-interactions of the synaptic PM localized NEGR1 result in an increase in this pool (Fig. 11). This increase correlates with a reduction in numbers of non-synaptic GAD65 clusters, indicating that NEGR1 promotes the recruitment of non-synaptic GAD65 to the synaptic PM. Fusion of synaptic vesicles with the PM may also result in the PM targeting of GAD65 attached to the synaptic vesicle membranes. We show, however, that the pool of GAD65 at the synaptic PM is not reduced and is even increased in tissues treated with tetanus toxin, which blocks fusion of synaptic vesicles with the PM. These experiments, electron microscopy data from previous studies showing GAD65 on some but not all synaptic vesicles, and biochemical studies showing that GAD65 does not co-purify with synaptic vesicles (Reetz et al., 1991; Takamori et al., 2006; Takamori et al., 2000) collectively indicate that GAD65 disassociates from vesicles before they fuse with the PM. Interestingly, synaptic vesicles contain high levels of palmitoyl protein thioesterase-1, an enzyme which de-palmitoylates GAD65 (Kim et al., 2008) and can therefore reduce the palmitoylation-dependent association of GAD65 with the vesicles (Kanaani et al., 2008). Deficiency in palmitoyl protein thioesterase-1 leads to a reduction in GAD65 levels in the brain (Kim et al., 2008) indicating that GAD65 de-palmitoylation is required for the maintenance of inhibitory synapses and GAD65 pool.

**Figure 11.**
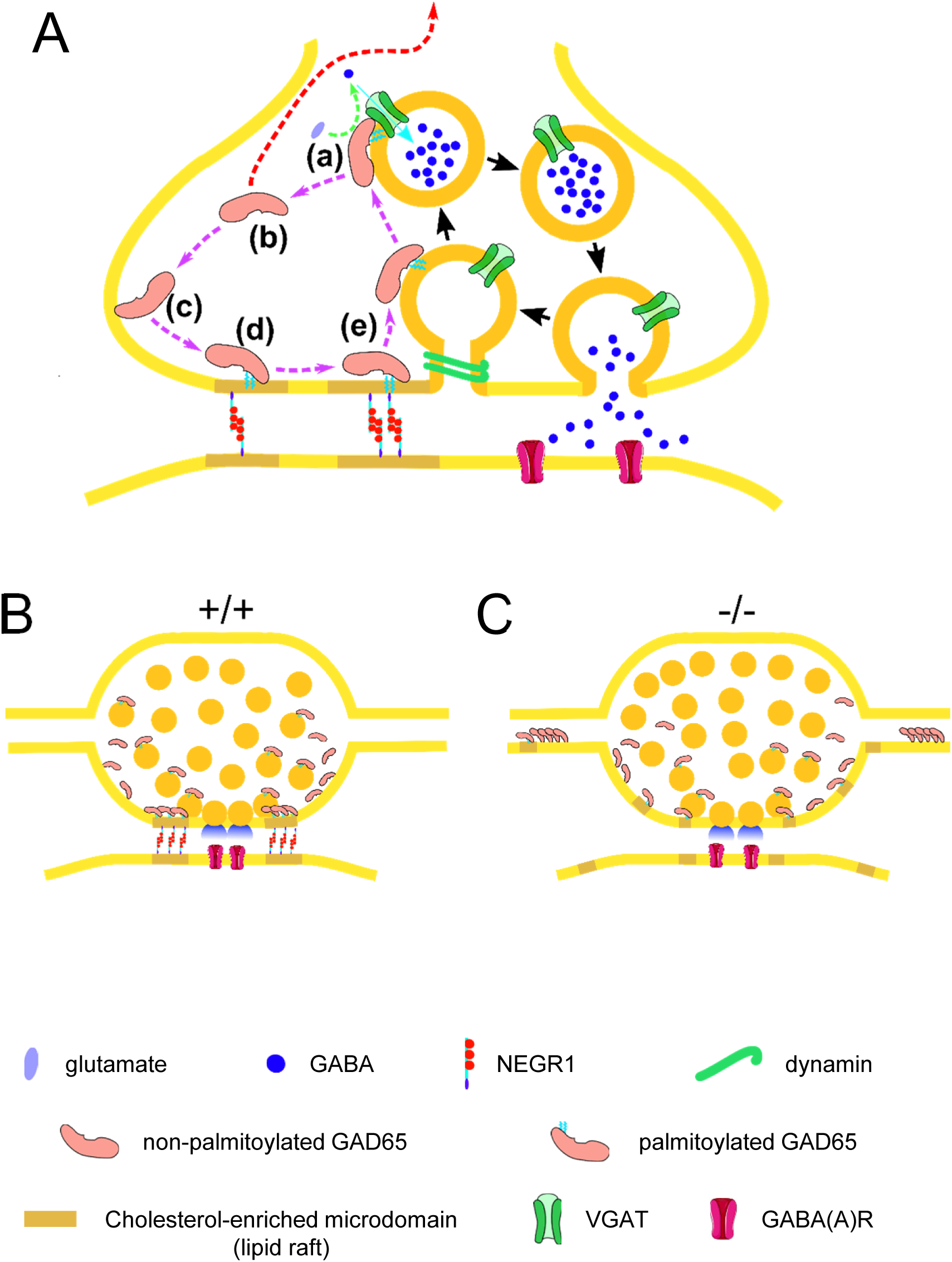
A hypothetical model describing NEGR1-dependent synaptic recruitment of GAD65. **(A)** Synaptic vesicle localized GAD65 uses glutamate to synthesize GABA, which is transported to the vesicle lumen by VGAT (**a**). GAD65 is de-palmitoylated and removed to the cytosol from refilled vesicles (**b**), which then fuse with the PM and release GABA. GAD65 released to the cytosol can travel to the Golgi in the neuronal cell body (red arrow, (Kanaani et al., 2008)) or attach to the PM in the synapse (**c**). Palmitoylation of GAD65 at the PM targets it to the NEGR1-containing cholesterol-enriched microdomains, which are anchored in synapses by NEGR1-containing adhesive bonds (**d**). NEGR1-dependent synaptic clustering of lipid rafts promotes ‘loading’ of palmitoylated GAD65 on the newly formed synaptic vesicles (**e**). This loading is facilitated by interactions between NEGR1 containing lipid rafts and components of synaptic vesicles constraining the retrieval of synaptic vesicle membranes. Synaptic recycling of vesicles and GAD65 is shown with black and pink arrows, respectively. **(B)** In NEGR1+/+ neurons, synaptic clustering of lipid rafts by trans-synaptic NEGR1-containing adhesive bonds promotes synaptic accumulation of GAD65. **(C)** In NEGR1-/- neurons, synaptic clustering of lipid rafts is reduced, synaptic targeting of GAD65 is inhibited, and non-synaptic GAD65 clusters are formed.

Cell adhesion molecules are involved in organizing the presynaptic nanoarchitecture required for highly regulated neurotransmitter release (Tang et al., 2016). NEGR1 is attached to the PM via a GPI anchor and accumulates in lipid microdomains, where palmitoylated proteins, including GAD65 (Kanaani et al., 2002) are also targeted. Trans-synaptic adhesive bonds formed by NEGR1 can promote the synaptic recruitment of GAD65 by limiting diffusion of lipid microdomains that GAD65 binds to. This scenario is supported by our observations showing that the synaptic clustering of lipid rafts is reduced in NEGR1-/- neurons. NEGR1-dependent synaptic clustering of lipid rafts can also constrain the retrieval of synaptic vesicle membranes, since synaptic vesicle proteins interact with PM-localized cholesterol (Thiele et al., 2000). Clustering of lipid rafts by GPI-anchored proteins triggers the assembly of the submembrane spectrin cytoskeleton (Leshchyns’ka et al., 2003) which also inhibits the retrieval of vesicle membranes (Puchkov et al., 2011). Our experiments showing that inhibition of synaptic endocytosis triggers clustering of GAD65 at the synaptic PM indicate that GAD65 is removed from the PM with membranes of reformed synaptic vesicles. An NEGR1-dependent restraint of vesicle reformation from the PM may be necessary for “loading” of GAD65 on the membranes of reforming synaptic vesicles, which may be facilitated within the cholesterol-enriched environment required for synaptic vesicle reformation (Martin, 2000), created by interactions between cholesterol enriched NEGR1-containing lipid microdomains and cholesterol-enriched synaptic vesicle membranes (Fig. 11). Palmitoylation plays a critical role in synaptic sorting of GAD65 (Kanaani et al., 2004; Kanaani et al., 2008). The GAD65-palmitoylating enzyme huntingtin-interacting protein HIP14 (Huang et al., 2004) is found at the neuronal PM and in Golgi and sorting/recycling and late endosomal structures in neurons (Huang et al., 2004), but is not detectable in synaptic vesicle preparations (Takamori et al., 2006). Slower reformation of vesicles may therefore facilitate the HIP14-mediated palmitoylation-dependent attachment of GAD65 to the membranes of vesicles during their retrieval from the PM.

We demonstrate that NEGR1 is highly expressed in NPY positive hypothalamic neurons and accumulates in the inhibitory GABAergic synapses formed by these neurons in culture. NPY/AgRP-expressing neurons play a prominent role in promoting food intake by inhibiting anorexigenic proopiomelanocortin (POMC) cells (Dietrich and Horvath, 2013; Suyama and Yada, 2018). We show that NEGR1 deficiency causes a reduction in numbers of inhibitory synapses in the ARC, where NPY/AgRP positive neurons form synapses on the POMC neurons (Dietrich and Horvath, 2013; Suyama and Yada, 2018). This synapse loss can be caused by impaired synaptic targeting of GAD65 in NEGR1-/- mice, because GAD65 deficiency causes a reduction in the size of the synaptic vesicle pool (Tian et al., 1999). We cannot exclude, however, that NEGR1-mediated adhesive bonds are also required for formation or maintenance of the GABAergic synapses. It is noteworthy that NEGR1 deficiency causes a reduction in the density of spines in cortical neurons (Pischedda et al., 2014; Szczurkowska et al., 2018). In cultured hippocampal neurons, NEGR1 expression peaks at 14 days *in vitro*, and NEGR1 overexpression in twenty-one-day-old hippocampal neurons, i.e., when the levels of endogenous NEGR1 decline, results in an increase in synapse density (Hashimoto et al., 2008), also suggesting that NEGR1 promotes synapse stabilization. A reduction in the efficiency of the inhibitory synaptic input into the anorexigenic POMC cells could cause a reduction in food intake observed in mice with inactivated NEGR1 (Lee et al., 2012) and could also cause a reduction in body mass ((Lee et al., 2012), observed also in our colony: at 2-3 months of age, mean ± SEM: 26.38 ± 0.54 g NEGR1+/+ males vs 23.63 ± 0.71 NEGR1-/- males, *p = 0.006 Mann-Whitney test; 20.43 ± 0.27 g NEGR1+/+ females vs 19.02 ± 0.39 NEGR1-/- females, *p = 0.004 Mann-Whitney test). A similar mechanism may also contribute to a reduction in the body mass index in humans with mutations affecting NEGR1 expression (Antunez-Ortiz et al., 2017).

NPY/AgRP neurons also play a key role in coordinating the activity of hypothalamic hunger circuits with the activity of midbrain reward circuits (Alhadeff et al., 2019). Our results, demonstrating that NEGR1 deficiency results in blunting of the reinforcing effects of food, indicate that NEGR1 also functions in the brain reward pathways. Dysregulated brain reward pathways may be contributing to increased intake of palatable foods and ultimately obesity (Berthoud et al., 2011). Mutations in noncoding regulatory elements upstream of *NEGR1* containing binding sites for transcriptional repressors are found in people with severe early-onset obesity (Wheeler et al., 2013; Willer et al., 2009), suggesting that NEGR1 overexpression causes changes in feeding behavior, which ultimately lead to obesity. We report that the levels of NEGR1 and its synaptic enrichment are increased in brains of mice with high fat diet induced obesity confirming a previous study, which found increased NEGR1 mRNA levels in the hypothalamus of 15-week-old female mice fed a high fat diet (Lee et al., 2010). High fat diet induces the loss of synapses in multiple brain regions including the hypothalamus (Dietrich and Horvath, 2013; Horvath et al., 2010; Lizarbe et al., 2018), hippocampus (Lizarbe et al., 2018; Stranahan et al., 2008; Valladolid-Acebes et al., 2012), and cortex (Bocarsly et al., 2015; Lizarbe et al., 2018). Overexpression of NEGR1 in fourteen-day old hippocampal neurons, i.e. at the peak of endogenous NEGR1 expression in these neurons, leads to a reduction in synapse density (Hashimoto et al., 2008). We demonstrate that NEGR1 overexpression causes a reduction in the density of synapses formed on dendrites and axons of NEGR1-overexpressing hypothalamic neurons, indicating that NEGR1 overexpression can directly contribute to the high fat diet induced synapse loss, including the loss of inhibitory synapses reported previously (Valladolid-Acebes et al., 2012). NEGR1 overexpression causes a strong inhibition of synaptic vesicle membrane retrieval and can therefore affect synapse formation or maintenance by inhibiting synaptic vesicle biogenesis, which is required for synapse formation during neuronal development (Hannah et al., 1999) and maintenance of neurotransmitter release in mature synapses (Milosevic, 2018). Similarly, the loss of inhibitory synapses was found in dynamin 1, 3 double knock-out mice (Raimondi et al., 2011). Furthermore, we demonstrate that the high fat diet and trans-interactions of NEGR1 cause similar reductions in synaptic GABA concentrations and induce similar increases of GAD65 levels at the PM, suggesting that the high fat diet caused reduction in the concentration of GABA, observed at least in some brain regions including the frontal cortex and hippocampus of rats (Sandoval-Salazar et al., 2016) and the prefrontal cortex of mice (Hassan et al., 2018), can also be associated with NEGR1 overexpression.

NEGR1 downregulation in mice also results in impaired core behaviors related to autism spectrum disorders (Szczurkowska et al., 2018). In humans, deletions of the *NEGR1* gene have been found in patients with developmental co-ordination disorder, attention deficit / hyperactivity disorder, learning disability, delayed speech and language development, and dyslexia (Genovese et al., 2015; Tassano et al., 2015; Veerappa et al., 2013). The results of our work suggest that NEGR1-dependent dysregulation of the GABAergic system can also be a factor contributing to the development of these disorders.

## Materials and Methods

### Antibodies

Goat polyclonal antibodies against NEGR1 (AF5394) used for immunocytochemistry (IC: 1:100) and immunohistochemistry (IHC: 1:1000) were from R&D systems. Rabbit polyclonal antibodies against NEGR1 (ab101809) used for treatment of living neurons (1:150), rabbit polyclonal antibodies against ganglioside GM4 (ab23947, IC: 1:100, Dot Blot (DB): 1:8000) and mouse monoclonal [R24] to ganglioside GD3 (ab11779, DB: 1:100) were from Abcam. Goat polyclonal antibodies against NEGR1 (sc-137625, Western blot (WB): 1:250), goat polyclonal antibodies against synaptophysin (sc-7568, IC: 1:50), mouse monoclonal antibodies against synaptophysin (sc-17750, WB: 1:1000), mouse monoclonal antibodies against VGAT (sc-393373, WB: 1:250), goat polyclonal antibodies against tau (sc-1995, IC: 1:100), rabbit polyclonal antibodies against NPY (sc-28943, IC: 1:200) and goat polyclonal antibodies against NCAM1 (sc-1507, WB: 1:500) were from Santa Cruz Biotechnology. Mouse monoclonal antibodies against MAP2 (M4403, IC: 1:100) and mouse monoclonal antibodies against GAPDH (WB: 1:1000) were from Sigma. Mouse monoclonal antibodies against GAD65 (GAD-6 supernatant, IC: 1:20, IHC: 1:150, WB: 1:20) were from Developmental Studies Hybridoma Bank. Mouse monoclonal antibodies against PIP2 [2C11] were from Novus Biologicals (NB100-2751, DB: 1:1000). Guinea pig polyclonal antibodies against the cytoplasmic domain of VGAT (#131004, IC: 1:200; IHC: 1:900) and rabbit polyclonal antibodies against the lumenal domain of VGAT (#131103, IC: 1:200) were from Synaptic Systems. Non-immune rabbit immunoglobulins were from Sigma. Fluorochrome- and HRP-conjugated secondary antibodies used for detection were from Jackson ImmunoResearch.

### DNA constructs

DNA coding for full length HA-tagged human NEGR1 (NM_173808.2) was synthesized using the GeneArt® Gene Synthesis service (Life Technologies) and subcloned into the pcDNA3 vector. GFP-tagged GAD65WT and GAD65(C30,45A) plasmids were generated and characterized as described previously (Kanaani et al., 2004; Kanaani et al., 2002; Kanaani et al., 2008; Phelps et al., 2016). mCherry-D4H encodes mCherry-tagged fourth domain of perfringolysin O theta toxin with D434S mutation (Maekawa and Fairn, 2015). To generate LCK-VN, DNA encoding amino acids 1-172 of Venus (VN) was amplified by PCR and cloned in frame after the first 26 amino acids of human LCK with a flexible glycine-serine linker (GSGSGSSGGGSSGSS). To generate GAD65-VC constructs, DNA encoding wild type and palmitoylation deficient GAD65 was PCR amplified from GAD65-GFP and GAD65-(30,45A)-GFP and cloned in frame with DNA coding for amino acids 158-238 of Venus (VC) using the same linker (GSGSGSSGGGSSGSS). GAD65-(24-31A)-VC was generated from GAD65-VC by site-directed mutagenesis using primers designed to mutate amino acids 24-31 of GAD65 to alanine. All constructs were cloned via AQUA cloning (Beyer et al., 2015) and verified by sequencing.

### Mice

C57Bl/6NJ mice for high fat diet experiments were obtained from the Australian Phenomics Network (APN) based at the Monash University, Melbourne, Australia, and housed at 22 ± 1 °C with a controlled 12-h light/dark cycle and had ad libitum access to water. Mice (9-11-week-old) were fed with either chow (8% calories from fat) or high-fat diets (HFD) (45% calories from fat) (Turner et al., 2009) for 8 weeks (UNSW animal care and ethics committee (ACEC) permit 17/120B). NEGR1+/+, NEGR1+/- and NEGR1-/- littermates (Negr1tm1b(KOMP)Wtsi mice, UCDAVIS KOMP Repository) were used for biochemical (2-3-month-old), behavioral (7-8-month-old) and immunohistological (7-8-month-old) analysis (UNSW ACEC permits 19/37B, 20/93B). Similar results were obtained for males and females and the data was combined. Newborn NEGR1+/+ and NEGR1-/- mice from homozygous breeding pairs were used for neuronal culture preparation (UNSW ACEC permits 17/102A, 20/93B). C57Bl/6 mice (2-3-month-old) were used for biochemical and neuronal culture experiments that did not analyze effects of NEGR1 deficiency (UNSW ACEC permit 18/99A). All experiments followed guidelines issued by the National Health and Medical Research Council of Australia.

### Production, purification and application of recombinant soluble NEGR1

DNA coding for soluble human NEGR1 (sNEGR1, amino acids 1-315 of NEGR1 above glycine 324 attached to the GPI anchor) was amplified using forward 5’- CACCATGGACATGATGCTGTTGGTGCA-3’ and reverse 5’-TGGAGGGTTAAGAGGCAGG-3’ primers, cloned into pEF5/FRT/V5 directional expression vector (Thermo Fisher Scientific) and transfected into CHO cells. Cells stably transfected with the construct were selected with Hygromycin B (Thermo Fisher Scientific) and maintained in culture in Ham’s F-12 culture medium (PAA Laboratories) with 5% foetal bovine serum (Thermo Fisher Scientific). Cell culture media containing sNEGR1 secreted by the cells was collected and centrifuged for 15 min at 5000 g at 4°C to remove cell debris. The protein was concentrated with 40% ammonia sulphate, desalted and purified using a continuous-elution electrophoresis cell (Model 491 Prep Cell, Bio-Rad) according to the manufacturer’s instructions as described (Leshchyns’ka et al., 2015). The purity of the protein was controlled by SDS-PAGE electrophoresis with subsequent silver staining and Western blot analysis.

### Treatment of hypothalamic tissue slices

Hypothalami were dissected, cut into 1 mm thick slices, placed into 1.5 ml tubes and incubated with recombinant sNEGR1 (10 µg/ml) or bovine serum albumin (10 µg/ml) diluted in Neurobasal A supplemented with gentamycin/amphotericin (1:500, ThermoFisher Scientific) for 24 h at room temperature with constant agitation. Tetanus toxin (Sigma, T3194, 1 nM) and dynasore (Tocris, Cat. No. 2897, 80 µM) were included when indicated. Tissues were collected by centrifuging tubes at 3000 g for 10 min at 4°C.

### Preparation of brain tissue homogenates, soluble protein fractions and synaptosomes

Homogenates (10%, w/v) were prepared in HOMO-A buffer (HOMO buffer (1 mM MgCl_2_, 1 mM CaCl_2_, 1 mM NaHCO_3_, 5 mM Tris, pH 7.4) containing 0.32 M sucrose, EDTA-free complete inhibitors (Roche) and 1 mM PMSF (Sigma)). Homogenates were used for synaptosome isolation as described (Andreyeva et al., 2010; Shetty et al., 2013). All steps were performed at 4°C. Briefly, homogenates were centrifuged at 1400 g for 10 min. The supernatant and pellet were resuspended in HOMO-A buffer and centrifuged for 10 min at 700 g. The resulting supernatants were combined and centrifuged at 17500 g for 15 min. The 17500 g supernatant was centrifuged at 200000 g for 1 h, the supernatant collected and used as the soluble protein fraction. The 17500 g pellet was resuspended in HOMO-A buffer and applied on top of a step gradient with interfaces of 0.65 M, 0.85 M, 1 M, 1.2 M sucrose in HOMO buffer. The 700 g pellets were combined, adjusted to 1 M sucrose in HOMO buffer and layered on 1.2 M sucrose in HOMO buffer. HOMO-A buffer was applied on top of the gradient. The crude synaptosomal fractions were collected at the 1 M / 1.2 M interface after centrifugation for 2 h at 100000 g and combined. The crude synaptosomal fraction was again adjusted to 1 M sucrose and layered on top of the 1.2 M sucrose. HOMO-A buffer was applied on top of the gradient. After centrifugation for 2 h at 100000 g, synaptosomes were collected at the 1 M / 1.2 M interface, resuspended in HOMO-A buffer, pulled down by centrifugation for 30 min at 100000 g and resuspended in HOMO-A buffer.

### Analysis of the GABA levels in synaptosomes

Serially diluted GABA standards (0 to 5 µM) were used to construct a standard calibration curve. All standards and samples contained a fixed amount of deuterium-labeled GABA internal standard (d6_GABA) to correct for variations during extraction and instrumental analysis. Synaptosomes were pelleted down by centrifuging them at 100000 g for 45 min at +4°C. The pellets were immediately frozen and stored at -80°C. Synaptosomes were extracted by adding 50 µl of water to tubes with synaptosomes followed by brief sonication and addition of 100 µl acetonitrile with vigorous vortexing. Internal standard (100 µl in acetonitrile) was added and the tubes were vortexed and incubated at - 20°C for 10 min. Tubes were centrifuged at 13000 rpm for 10 min and pellets containing proteins, buffer salts and sugars were discarded. Supernatants were dried, reconstituted in 100 µl acetonitrile: water (4:1) and filtered through 4 mm x 0.2 µm disks (Phenomenex, Australia) into LC vials. For highly specific and sensitive measurement of GABA, analysis was carried out on a LC–MS/MS system consisting of Accela Open AS autosampler, Accela UPLC pump coupled to a Quantum Access triple quadrupole mass spectrometer equipped with a heated electrospray (HESI) probe (Thermo Scientific, San Jose, CA, USA). The capillary and spray temperatures were both set to 300°C and the electrospray capillary voltage to 4 kV. The dwell time for each transition was set to 250 ms. Argon was used as the collision gas at a pressure of 1 torr. Standards and synaptosome extracts (20 µl each) were injected onto a Luna NH₂ column (3 µm, 2.0 mm × 150 mm) (Phenomenex, Australia) maintained at room temperature. The flow rate was set at 300 µl / min. Mobile phase A was a mixture of 10% aqueous 5 mM ammonium acetate at pH 9 and 90% neat acetonitrile. Mobile phase B was 100% aqueous 5 mM ammonium acetate at pH 9. A gradient program was used for the optimum retention of GABA followed by column clean up and equilibration. The percentage of mobile phase B increased linearly according to the gradient: 0 min, 20%; 2 min, 20%; 0.5 min, 40%; 2 min, 60%; held 3 min, 20%; 0.5 min, 20% held 7 min. Acquisition of mass spectrometric data was performed in selected reaction monitoring mode (SRM) in positive heated electrospray ionisation (HESI) using the transitions of m/z 104.2 --> 87.2 for GABA and m/z 110.2 --> 93.2 for d6-GABA. Data analysis was performed using Thermo Scientific Xcaliburᵀᴹ software (version 2.07), with the LCQuan feature of this software used for automated data analysis. Software generated peak area ratios (GABA/d6-GABA) were plotted against standard concentrations to construct the calibration curve (r² = 0.995). The endogenous concentration of GABA in samples was extrapolated from this curve based on the area ratios obtained for each sample. The final concentration was normalized to protein values.

### Analysis of GABA content from INS-1E cells

INS-1E cells were seeded at a density of 15,000 cells / cm^2^. After 24 h of culture, cells were transfected using Lipofectamine 2000 with plasmids containing either GAD65-GFP or GAD65(C30,45A)-GFP mutant. Three days post-transfection, cells were collected and lysed directly in mobile phase (3% methanol, 17% acetonitrile in 0.1M phosphate buffer at pH 6). One part lysate was diluted with two parts methanol and kept at -20°C for 20 min prior to filtering (Millipore Ultrafree-MC, 0.22 μm). GABA and glutamate content were analyzed using electrochemical detection of OPA-derivatized amino acids. Reverse phase HPLC was used to separate amino acids (EICOM HTEC-510 ECD with FAO3DS column) prior to electrochemical detection (Zandy et al., 2017). Serial dilutions of GABA and glutamate were analyzed to generate a linear calibration curve using area under the curve analysis from chromatograms.

### Isolation of synaptic PM

Synaptic PM was isolated from synaptosomes as described (Leshchyns’ka et al., 2006). Unless otherwise stated, all steps were performed at 4°C. Briefly, synaptosomes were lysed by diluting them in 9 volumes of ice-cold H_2_O and then immediately adjusted by 1 M HEPES, pH 7.4 to a final concentration of 7.5 mM HEPES. After incubation on ice for 30 min, the mixture was centrifuged at 100000 g for 20 min and the pellet containing synaptic PM was collected.

### SDS-PAGE electrophoresis and Western blot analysis

Proteins were separated on a 4-12% Bis-Tris bolt mini gel (Life Technologies) and electroblotted onto Hybond PVDF transfer membrane (0.2 µm; Amersham). Molecular weight markers were pre-stained protein standards from Bio-Rad. Membranes were washed with PBS-Tween, blocked for 1 h at room temperature with 5% w/v skim milk in PBS-Tween, incubated with appropriate primary antibodies overnight, washed with PBS-Tween and incubated with corresponding HRP-conjugated secondary antibodies for 1 h at room temperature. Luminata Forte Western HRP substrate (Merck Millipore) or Western Lightning Ultra (Perkin Elmer) was applied onto membranes to visualize labelling. Images were captured using Micro-Chemi 4.2 (DNR Bio-Imaging Systems) or ChemiDoc XRS+ Imaging System (Bio-Rad).

### Culture, transfection and treatment of primary hypothalamic neurons

Primary hypothalamic neurons were prepared by dissecting hypothalami from brains of 1-3-day-old mice and disassociating neurons as described (Sheng et al., 2015; Sheng et al., 2019). Neurons were maintained in Neurobasal A medium supplemented with 2% B-27, GlutaMAX and FGF-2 (2 ng/ml) (all reagents from Thermo Fisher Scientific) in 24-well plates for 2 weeks on coverslips coated with poly-D-lysine (100 µg/ml). Neurons were transfected before plating by electroporation using a Neon transfection system (Thermo Fisher Scientific). Alternatively, neurons were transfected using the calcium phosphate method essentially as described (Jiang and Chen, 2006). Briefly, coverslips with neurons maintained for 7 to 10 days in culture were transferred to new wells of a 24-well plate containing 500 µl of growth medium 30 min prior to transfection. One µg of plasmid DNA and 3.1 µl of 2 M CaCl_2_ were mixed with water to a final volume of 25 µl per coverslip. DNA-Ca^2+^-phosphate precipitate was prepared by adding DNA/CaCl_2_ solution to 25 µl of 2x HEPES-buffered saline (HBS, 280 mM NaCl, 1.5 mM Na_2_HPO_4_, 50 mM HEPES, pH 7.06) (1/8^th^ at a time and mixing briefly between each addition). The solution was incubated for 10 min at 37°C and the resulting suspension was applied dropwise to the coverslips. Neurons were incubated with the precipitate for 3 h, washed with acidified Tyrode’s solution (pH 6.7-6.8) to remove the precipitate, and transferred back to the wells containing the original conditioned culture medium. When indicated, soluble recombinant NEGR1_1-315_ (10 µg/ml prepared using a stock solution in water), tetanus toxin (1 nM prepared using a stock solution in water) or dynasore (80 µM prepared using a stock solution in DMSO) were applied to neurons in the culture medium. Control neurons were treated with the culture medium containing the same concentration of the vehicle used to prepare the stock solutions.

### Labelling of live hypothalamic neurons with antibodies against the lumenal domain of VGAT

Neurons were pre-incubated for 10 min with sNEGR1 (10 µg/ml) in the culture medium or mock treated with the culture medium at 37°C and 5% CO_2_. Rabbit polyclonal antibodies against the lumenal domain of VGAT were then added to live neurons and incubated for 30 min at 37°C and 5% CO_2_. Further steps were performed at room temperature. Neurons were gently fixed with 4% paraformaldehyde in PBS for 15 min, washed with PBS, blocked with 1% donkey serum in PBS for 15 min, washed with PBS, and labelled with anti-rabbit Cy3-conjugated secondary antibodies in 1% donkey serum in PBS for 30 min to visualize the antibodies against the lumenal domain of VGAT remaining at the cell surface. Neurons were then washed with PBS, post-fixed with 2% formaldehyde in PBS for 5 min, permeabilized with 0.25% Triton X-100 in PBS for 5 min, blocked with 5% donkey serum in PBS for 20 min and incubated with guinea pig polyclonal antibodies against the cytoplasmic domain of VGAT applied in 1% donkey serum in PBS overnight at +4°C. Neurons were then washed with PBS and labelled with a cocktail of anti-rabbit Cy5-conjugated and anti-guinea pig Cy2-conjugated secondary antibodies in 1% donkey serum in PBS for 30 min to visualize the total pool of antibodies against the lumenal domain of VGAT and the total pool of synaptic VGAT, respectively. Immunofluorescence images were captured at room temperature using a confocal laser scanning microscope C1si (Nikon, Tokyo, Japan), NIS-elements software (version 4.0; Nikon), and Plan Apo VC 60x oil-based objective (Nikon, numerical aperture 1.4). Accumulations of synaptic VGAT detected with antibodies against its cytoplasmic domain were outlined using a threshold function of ImageJ (National Institute of Health, Bethesda, MD), and labelling intensities of the cell surface and total pool of antibodies against the lumenal domain of VGAT were measured within the outlines.

### Immunofluorescence labelling and analysis of fixed neurons

The labelling was performed essentially as described previously (Bliim et al., 2019; Sytnyk et al., 2002). Unless stated otherwise, all steps were performed at room temperature. Neurons were fixed with 4% formaldehyde in PBS for 15 min, washed with PBS, permeabilized with 0.25% Triton X-100 in PBS for 5 min, blocked in 5% donkey serum in PBS for 20 min, and incubated with primary antibodies applied in 1% donkey serum in PBS overnight at +4°C. Neurons were then washed with PBS, incubated for 45 min with corresponding secondary antibodies in 1% donkey serum in PBS, washed with PBS and embedded in the ProLong Gold Antifade mounting medium (Thermo Fisher Scientific). Immunofluorescence images were captured at room temperature using a confocal laser scanning microscope C1si (Nikon, Tokyo, Japan), NIS-elements software (version 4.0; Nikon), and Plan Apo VC 60x oil-based objective (Nikon, numerical aperture 1.4). To measure labeling intensities of synaptophysin along axons and dendrites, 50 – 100 µm long fragments of proximal dendrites and all fragments of axons (over 30 µm long) located within 40 µm from the soma were manually outlined in ImageJ, and mean labeling intensities were measured within the outlines. To measure labeling intensities of GAD65 in synaptophysin accumulations, they were outlined using a threshold function of ImageJ, and GAD65 labelling intensities were measured within the outlines. To calculate percentages of synaptic and non-synaptic GAD65 clusters, all GAD65 accumulations were outlined using a threshold function in ImageJ, and the presence of synaptophysin labelling within the outlines was then manually analyzed.

### Analysis of the activity-dependent FM4-64 dye uptake and release in neurons

The analysis was performed essentially as described previously (Andreyeva et al., 2010; Leshchyns’ka et al., 2006; Shetty et al., 2013). When indicated, neurons were incubated for 10 min at room temperature with rabbit polyclonal antibodies against NEGR1 or non-immune rabbit immunoglobulins diluted in 4 mM K^+^ buffer (150 mM NaCl, 4 mM KCl, 2 mM MgCl_2_, 10 mM glucose, 10 mM HEPES, 2 mM CaCl_2_). To load synaptic vesicles with FM4-64, neurons were incubated with FM4-64 (15 μM, Thermo Fisher Scientific) diluted in 47 mM K^+^ buffer (107 mM NaCl, 47 mM KCl, 2 mM MgCl_2_, 10 mM glucose, 10 mM HEPES, 2 mM CaCl_2_) for 2 min at room temperature and then washed with 4 mM K^+^ buffer for 10 min. Images of neurons were taken using a Nikon Eclipse Ti microscope equipped with the CoolLED light source, Nikon DS-Qi1Mc or Andor Zyla 4.2+ sCMOS camera and 60x Plan Apo VC water immersion objective (Nikon, numerical aperture 1.2) using 550 nm excitation wavelength. In experiments with transfected neurons, GFP fluorescence detected using 488 nm excitation wavelength was used to identify dendrites and axons of transfected hypothalamic neurons. Time lapse images were acquired with a 1 s interval before (10 baseline images) and during electric field stimulation (1 ms bipolar pulses at 10 Hz for 100 s) applied using S88X stimulator (Grass Technologies). FM4-64 uptake was analyzed by outlining accumulations of the dye with the mean intensity at least 2-fold higher than the background and measuring the mean intensity of the labeling within the outlines in ImageJ. To analyze FM4-64 dye release, the mean intensity of FM4-64 labelling within the outlines was measured over time. Background fluorescence intensity was measured in the FM4-64-free area of the image and subtracted from the data collected in FM4-64 accumulations. The half-life of FM4-64 fluorescence decay in each bouton was measured using a non-linear one-phase exponential decay analysis in Prism7. FM4-64 accumulations which did not show a stimulation-induced exponential decay in the intensity were excluded from the analysis. These accumulations typically had a very low amplitude of FM4-64 release and most likely represented endosomes.

### CHO cell culture, transfection and FM4-64FX dye loading

CHO cells were cultured in 24-well-plates on poly-D-lysine coated glass coverslips in the growth medium (Dulbecco’s Modified Eagle’s Medium / Nutrient Mixture F-12 HAM (DMEM/F12, D8437; Sigma) supplemented with 5% foetal bovine serum (Sigma)) in an incubator at 37°C and 5% CO_2_. Cells were transfected using calcium phosphate method. Thirty min prior to transfection, growth medium was replaced with DMEM/F12. For each coverslip, 1 µg of plasmid DNA and 3.1 µl of 2 M CaCl_2_ were mixed with water to a final volume of 25 µl. DNA-Ca^2+^-phosphate precipitate was prepared by adding DNA/CaCl_2_ solution to 25 µl of 2x HEPES-buffered saline (HBS, 280 mM NaCl, 1.5 mM Na_2_HPO_4_, 50 mM HEPES, pH 7.06) (1/8^th^ at a time and mixing briefly between each addition) and incubating for 10 min at 37°C. The resulting suspension was applied dropwise to the coverslips and incubated at 37°C and 5% CO_2_ for 3 h. Cells were then treated with 15% glycerol in DMEM/F12 for 1–2 min in the incubator, washed with PBS, placed in growth medium, and maintained for 24 h at 37°C and 5% CO_2_, and then used for analysis. When indicated, CHO cells were loaded with FM4-64FX dye (15 μM, Thermo Fisher Scientific) applied for 15 min at 37°C and 5% CO_2_ in 4 mM K^+^ buffer, washed 3 times with PBS, fixed and embedded in the ProLong Gold Antifade mounting medium (Thermo Fisher Scientific).

### Operant Conditioning Task

Prior to start of the experiment, mice were put on diet restriction for 7-15 days to achieve the weight loss to 80-85% of their original weight. They were kept on diet restriction throughout the experiment to maintain the weight of mice at 80-85% of their original weight. During the experiments, the mice were placed into an operant chamber containing two levers. The left “active” lever was connected to sucrose pellet dispenser releasing sucrose pellets into a food-magazine. The right lever was inactive. An operant conditioning test was conducted in three stages consisting of magazine training, fixed ratio training and progressive ratio test. The magazine training lasted for 30 min during which palatable sucrose pellets were automatically delivered from the food-dispenser to the food-magazine at a rate of 1 pellet / min. The numbers of times mice entered the magazine and numbers of pellets eaten by mice were counted. The magazine training was conducted for 6 to 10 days until the mouse left 13 or less pellets in the magazine by the end of the training session. The mice were then trained to press the lever in the fixed ratio training. During this training, active lever press but not inactive lever press resulted in the delivery of a single pellet into the food magazine. Each fixed ratio training session lasted for ≤ 30 min or until 30 pellets were delivered. Fixed ratio training was performed for ten days until the mouse left <15 pellets in the food-magazine by the end of the session. The mice were then subjected to the progressive ratio test essentially as described (June and Gilpin, 2010) during which the number of active lever presses was progressively increased to activate the release of the pellet. The number of presses required to release each consecutive pellet was calculated as described by Richardson and Roberts (1996) (Sharma et al., 2012) using the following formula: [5e ^(R*0.2)^] – 5, where R denotes the number of pellets rewarded already plus 1. Each PR session lasted for 1 h or until the break point at which mice refused to press the lever the number of times required to obtain the next pellet. The total number of times the active lever was pressed and the number of magazine entries were also counted as described (June and Gilpin, 2010). The progressive ratio sessions were performed on three consecutive days.

### Immunohistochemical analysis

Mice were deeply anaesthetized with intraperitoneally injected pentobarbitone sodium (100 mg / kg) and transcardially perfused with ice-cold PBS for 2 min followed by ice-cold 4% paraformaldehyde in PBS for 5 min. Brains were extracted and post-fixed in 4% paraformaldehyde for 24 h at 4°C, cryoprotected by incubation in 10% sucrose in PBS for 24 h and then stored in 30% sucrose in PBS at 4°C. Brains were cut coronally into 40 µm thick slices using a cryostat (Leica CM 1950) at -20°C. The slices were transferred into wells of 24 well plates containing PBS with 0.02% sodium azide and stored at 4°C. Two brain slices per animal corresponding to the same region of the hypothalamus and hippocampus (interaural 2.1 mm, bregma -1.7 mm) were selected. These slices were transferred into a new 24 well plate. Unless indicated otherwise, all further incubations were done on a shaker at room temperature. Slices were washed 3 times with PBS, and treated with sodium citrate buffer (pH 6.0, 10 mM) for 10 min at 70°C in a water bath to retrieve antigens. Slices were cooled down to room temperature, washed 3 times with PBS, blocked with 1% donkey serum and 0.25% Triton X-100 in PBS for 1 h, and incubated with primary antibodies diluted in 1% donkey serum in PBS overnight at 4°C, and then washed 3 times with PBS. Further steps were performed in dark. Slices were incubated with fluorochrome-conjugated secondary antibodies diluted in 1% donkey serum in PBS for 2 h, washed 3 times in PBS, counterstained with Hoechst dye in PBS for 10 min, washed with PBS, transferred to microscope slides with the help of a paintbrush and embedded in Prolong Diamond Antifade mounting medium. Immunofluorescence images were taken with a confocal laser scanning microscope C1si (Nikon, Tokyo, Japan), NIS-elements software (version 4.0; Nikon), and CFI Plan Apochromat VC 60XH objective (Nikon, numerical aperture 1.4).

## Acknowledgments

The study was supported by the National Health and Medical Research Council (VS), Australian Research Council (VS, HY), University of New South Wales (IL, VS), Australian Postgraduate Award (RPAT), University of New South Wales International Postgraduate Award (FS) and Tuition Fee Scolarship (IK), and Australian Government Research Training Program scholarship (SS, RK). The funders had no role in study design, data collection and analysis, decision to publish, or preparation of the manuscript.

## Competing interests

The authors declare that they have no competing interests with the contents of this article.

## Notes

### Competing Interest Statement

The authors have declared no competing interest.

